# The 100-plus Study of cognitively healthy centenarians: rationale, design and cohort description

**DOI:** 10.1101/295287

**Authors:** Henne Holstege, Nina Beker, Tjitske Dijkstra, Karlijn Pieterse, Elizabeth Wemmenhove, Kimja Schouten, Linette Thiessens, Debbie Horsten, Sterre Rechtuijt, Sietske Sikkes, Frans W.A. van Poppel, Hanne Meijers-Heijboer, Marc Hulsman, Philip Scheltens

## Abstract

**RATIONALE:** Although the incidence of dementia increases exponentially with age, some individuals reach >100 years with fully retained cognitive abilities. To identify the characteristics associated with the escape or delay of cognitive decline, we initiated the 100-plus Study (www.100plus.nl).

**DESIGN:** The 100-plus Study is an on-going prospective cohort study of Dutch centenarians who self-reported to be cognitively healthy, their first-degree family members and their respective partners. We collect demographics, life history, medical history, genealogy, neuropsychological data and blood samples. Centenarians are followed annually until death. PET-MRI scans and feces donation are optional. Almost 30% of the centenarians agreed to post-mortem brain donation.

**COHORT DESCRIPTION:** To date (September 2018), 332 centenarians were included in the study. We analyzed demographic statistics of the first 300 centenarians (25% males) included in the cohort. Centenarians came from higher socio-economic classes and had higher levels of education compared to their birth cohort; alcohol consumption of centenarians was similar, and most males smoked during their lifetime. At baseline, the centenarians had a median MMSE score of 25 points (IQR: 22.0-27.5); the large majority lived independently, retained hearing and vision abilities and was independently mobile. Mortality was associated with cognitive functioning: centenarians with a baseline MMSE score ≥26 and <26 points had a mortality percentage of respectively 17% and 42% per annual year in the second year after baseline (p=0.003). The cohort was 2.1-fold enriched with the neuroprotective *APOE-ε2* allele relative to 60-80 year-old population controls (p=4.8×10^-7^), *APOE-ε3* was unchanged and the *APOE-ε4* allele was 2.3-fold depleted (p=6.3×10^-7^).

**CONCLUSIONS:** Comprehensive characterization of the 100-plus cohort of cognitively healthy centenarians might reveal protective factors that explain the physiology of long-term preserved cognitive health.

## BACKGROUND

Although increasing age is the strongest risk indicator for cognitive decline and dementia, it is not an inevitable consequence of aging. The incidence of overall dementia starts to increase exponentially from approximately 60 years and at age 100 years the annual dementia incidence reaches 40% per year [1, 2]. However, the mere existence of cognitively healthy individuals older than 110 years [3-6] leads to the intriguing suggestion that the incidence of dementia decelerates somewhere after 100 years (See **Box**). Factors that allow for the preservation of cognitive health may thus be enriched for in super-agers, individuals who reach extreme ages with full cognitive functions [7]. The combination of extreme old age with maintained cognitive health is often observed in families [8-13], suggesting that beneficial factors involved in the long-term maintenance of both cognitive and overall health are heritable, and likely genetic [14, 15, 7, 16]. Indeed, results from the New England Centenarians Study indicated that siblings from centenarians are ∼ 8-12 times more likely to reach 100 years compared to individuals with no centenarian siblings [17].

This raises several questions: what are the unique molecular mechanisms that cause resilience against age related decline? Which hereditable factors are involved, and what is the role of the immune system? The answers to these questions are likely to provide novel insights in the effects of aging on the brain and they will be informative for the design of novel strategies that intervene in processes that lead to neurodegenerative diseases [18]. Answers to these questions might be found in the context of prospective follow-up studies, however, this is complicated by the fact that only ∼0.6% of the population born in the early 1900’s reaches 100 years (see **Box**). Therefore, we set out to identify protective factors in a cohort of centenarians who self reported to be cognitively healthy. For this, we initiated the 100-plus Study in 2013 at the Alzheimer center at the Amsterdam University Medical Center, Amsterdam, the Netherlands (www.100plus.nl). To date, the cohort includes 332 centenarians.

Children of centenarians also profit from the advantage they inherited by their centenarian parent: they live longer, and have almost 90% lower risk of developing myocardial infarction, stroke and diabetes compared to age-matched peers whose parents have average life spans [15, 19, 20]. Together, this suggests that first-degree family-members of centenarians are also enriched for protective (genetic) factors and that efforts to identify protective factors should include targeting the families of centenarians [21]. The value of using by-proxy phenotypes for genetic studies was recently demonstrated for 12 diseases [22], and recently for Alzheimer’s Disease [23, 24]. Application of the proxy-phenotype strategy to investigate extreme longevity offers additional value since using centenarian children as phenotype-by-proxy provides the opportunity to test relative to age-matched controls [25]. For this reason, we extended the 100-plus Study with a second phase in 2017, in which we also include first-degree family-members of centenarians and their partners.

The 100-plus Study has a main focus on the biomolecular aspect of preserved cognitive health. It is beneficial that cohort inclusion is on-going, as this allows us to take optimal advantage of the recent developments in high-throughput biomolecular techniques. For example, genetic variants of interest can be functionally tested in our collection of fresh blood samples and brain tissues from carriers.

Here we present the rationale for the 100-plus Study (See **Box**), we describe the study design and procedures, and we introduce the 100-plus Study cohort based on the clinical presentation of the centenarians at baseline, and the demographic characteristics of the centenarians relative to their birth cohort.

## METHODS/STUDY DESIGN

Please find in the **electronic supplementary material** (ESM.pdf) a complete compendium of participant inclusion procedures and current data collection procedures of the 100-plus Study.

### Inclusion and exclusion criteria

The 100-plus Study includes (i) Dutch-speaking centenarians who can (ii) provide official evidence for being aged 100 years or older, (iii) self-report to be cognitively healthy, which is confirmed by an informant (i.e. a child or close relation), (iv) consent to donation of a blood sample and (v) consent to (at least) two home-visits from a researcher, which includes an interview and neuropsychological testing. In the second phase of the 100-plus Study (from September 2017 onwards) we include (i) siblings or children from centenarians who participate in the 100-plus Study, or partners thereof who (ii) agree to donate a blood sample, (iii) agree to fill in a family history, lifestyle history and disease history questionnaire. Study exclusion criteria are limited to subjects who are legally incapable.

### Recruitment and research visits

#### Recruitment

We regularly perform an online search for local newspaper articles that mention a centenarian. These reports commonly include the name of the centenarian, and sometimes a description of their well-being and living situation. We retrieve an address online and we approach a prospective study participant by letter. When they express their interest in study participation and inclusion criteria are met, we schedule two baseline visits. (**See ESM.pdf** for detailed recruitment procedures).

#### Baseline visit

A researcher, trained to perform standardized visit procedures, will visit the centenarian. The baseline visit (T0) consists of two visits. The first baseline visit takes approximately 2 to 3 hours, and comprises obtaining informed consent for study inclusion, a life-history interview, an assessment of genealogy, and an assessment of current health and medical history **(Table 1)**. The second baseline visit, approximately one week after the first, takes approximately 1.5 hours: during this visit we subject the centenarian to a battery of neuropsychological tests and we measure grip strength and blood pressure **(Table 2)**. During the first baseline visit we inform participants of optional parts of the 100-plus Study: feces collection, PET-MRI or PET-CT imaging and post-mortem brain donation. Once a centenarian volunteers to participate in these parts of the 100-plus Study, we obtain informed consent for these study parts separately.

**Table 1:**
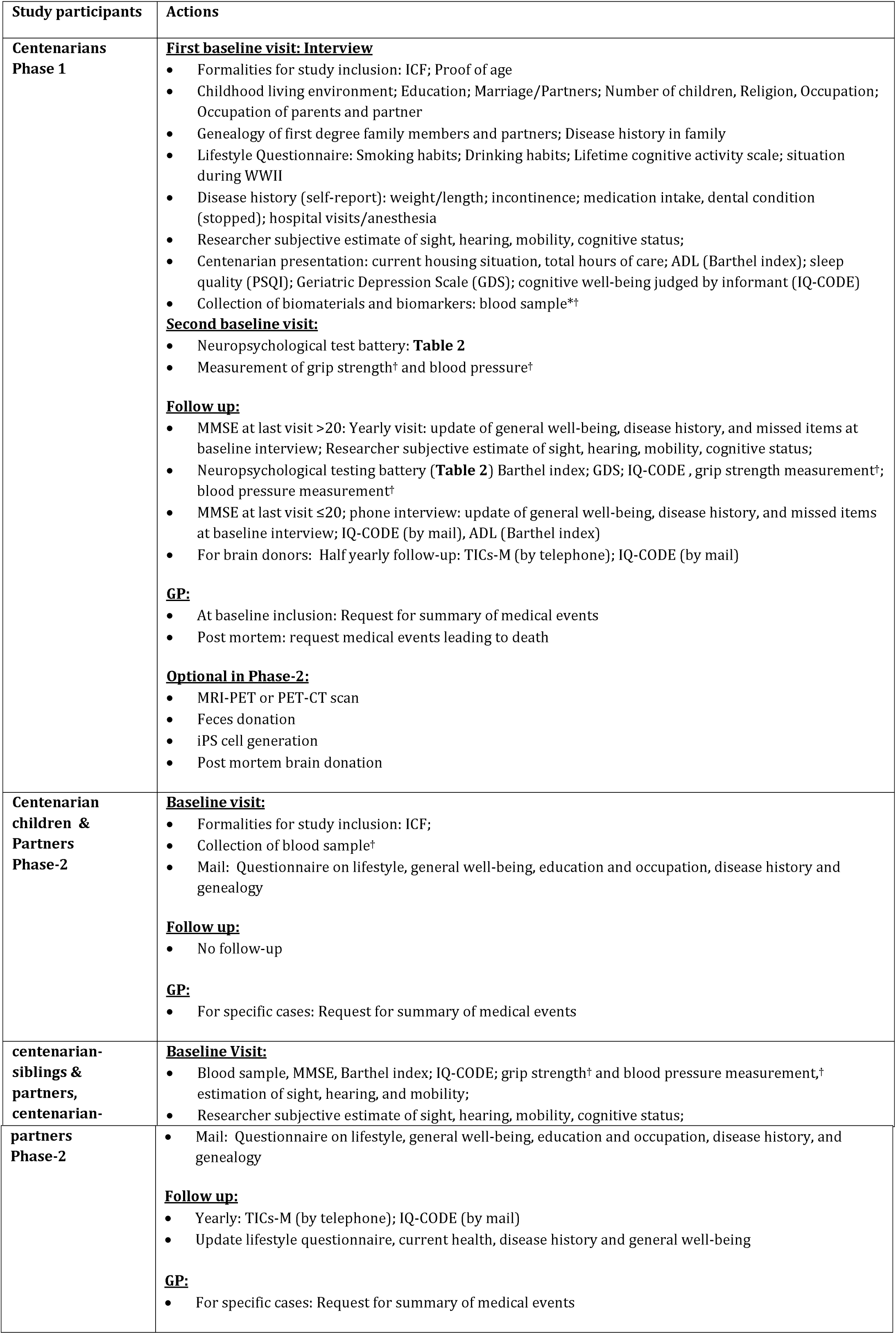
Overview of 100-plus Study data-collection. *Blood sample collection may occur at a different occasion, close to first baseline visit; Phase-2 of the 100-plus Study started in September 2017. †Blood sample biomarkers determined in the blood sample, assessment of blood pressure and measurement of grip strength are described in detail in ESM.pdf. TICS-M: Telephone Interview Cognitive Status –Modified (see Table 2); IQ-COde Informant Questionnaire on Cognitive Decline in the elderly short form (see Table 2).

**Table 2:** Neuropsychological tests and questionnaires. a: only administered at baseline; b: in 100-plus Study-phase 1 only. c: Included with the confirmation letter of study-inclusion, collected during the first baseline visit. d: Only administered during half yearly-follow up of brain donors and yearly follow-up of siblings

| Domain or goal | Assessment/Questionnaires | Duration (min) |
| --- | --- | --- |
| **Cognitive functioning** |  |  |
| Overall cognitive functioning | Researcher subjective impression of cognitive health (see methods)<br>Mini-Mental State Examination [26, 87]<br>National Adult Reading Test<sup>a</sup> [88-90]<br>Telephone Interview Cognitive Status –Modified (TICS-M)<sup>d</sup>[91] | 0<br>5<br>3 |
| Memory | CERAD 10-word list – immediate and delayed recall [92]<br>Visual Association Test - Memory [93]<br>Rivermead Behavioral Memory Test (RBMT)<sup>b</sup> immediate and delayed recall [94, 95] | 15<br>5<br>6 |
| Attention | Digit Span – forwards [96-98]<br>Trail Making Test A [99, 100] | 3 |
| Executive functions | Digit Span – backwards [96-98]<br>Letter Fluency – DAT [101-105]<br>BADS – subtest Key Search [106, 107]<br>BADS– subtest Rule Shift Cards [106, 107]<br>Trail Making Test B [99, 100]<br>Amsterdam Dementia Screening Test - Meander figure [108] | 3<br>2<br>3<br>3<br>10<br>2 |
| Language | Category Fluency – Animals [101, 109, 102]<br>Visual Association Test - Naming [93] | 2<br>1 |
| Visuo-spatial functioning/construction | CAMDEX-R/N CAMCOG - Figure Copying [110, 111]<br>Clock Drawing Test [112, 113]<br>Visual Object and Space Perception (VOSP) Battery<sup>b</sup> - subtest Number Location [114] | 3<br>2<br>3 |
| **Depression, ADL, Sleep, Lifestyle, geriatric impairments** |  |  |
| Depressive symptoms | Geriatric Depression Scale-15 (GDS) [29, 30] | 4 |
| (Instrumental) Activities of daily living | Informant Questionnaire on Cognitive Decline in the elderly short form (IQ-CODE) [31, 32]<br>Barthel Index [27, 115-117] | 3<br>3 |
| Lifetime cognitively stimulating experience | Lifetime Cognitive Activity Scale<sup>a</sup> [35, 36] | 5 |
| Sleep quality | Pittsburgh Sleep Quality Index<sup>a</sup> (PSQI) [28] | 5 |
| Geriatric impairments | Researcher subjective impression of sight, hearing, mobility | 0 |

#### Follow-up visits

During yearly follow-up visits (T1, T2…), which take approximately 2 hours: we inform about possible changes in cognitive functioning that took place in the last year, we update the interview questionnaire and re-administer the complete cognitive test battery and physical measurements **(Table 1, 2)**. Follow-up is continued until the participant is no longer willing/able to participate. When the MMSE score declines ≤20 there is evidence of clear cognitive impairment [26], and subjecting a centenarian to a neuropsychological testing battery becomes more complicated and follow-up visits by a researcher may no longer be constructive. When the MMSE at last visit drops below 20 (imputed MMSE score), we follow-up by informant questionnaire. To ensure up-to-date cognitive health measurements of brain donors, we administer telephone an informant questionnaires 6 months after the annual visit (T0.5, T1.5…). For a diagram of procedures see **Fig 2**. We ask informants to inform us when a participant dies and about the events that preceded death.

**Fig. 1:**
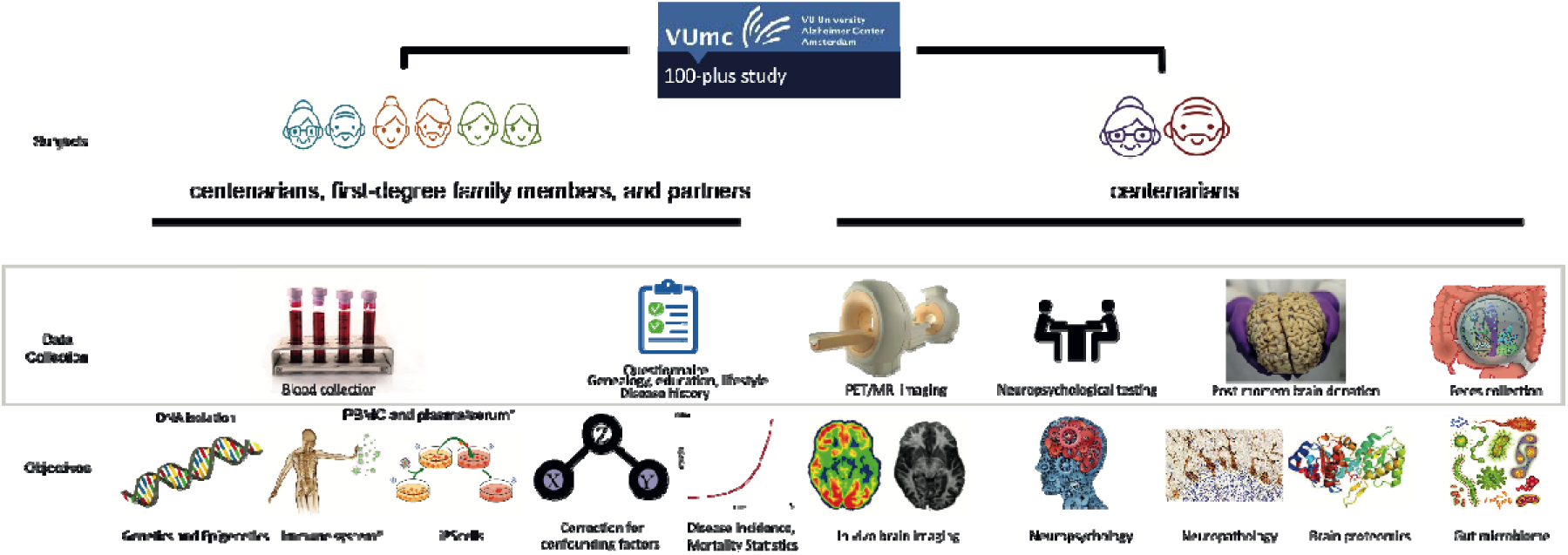
Overview of the 100-plus Study, Phase 2: During home visits we inquire about life-history of the centenarians, their family history, medical history, and current health. We assess their performance on neuropsychological tests, measure blood pressure and grip strength and we collect a blood sample, for blood testing and genetic analyses. Optional parts of the study are: a visit to the VUmc clinic for PET-MRI and/or PET-CT imaging, feces donation to investigate the gut microbiome, and the generation of iPS cells from peripheral blood. Furthermore, all participants are informed about the option of post-mortem brain donation in collaboration with the Netherlands Brain Bank [38]. This is optional and not required for study participation. We evaluate changes in general well-being and in neuropsychological test performance during (half-)yearly follow-up visits. Next to the centenarians, we also include their first-degree family members and their partners. * Collected in Phase-2 of the 100-plus Study, started in September 2017.

**Fig. 2:**
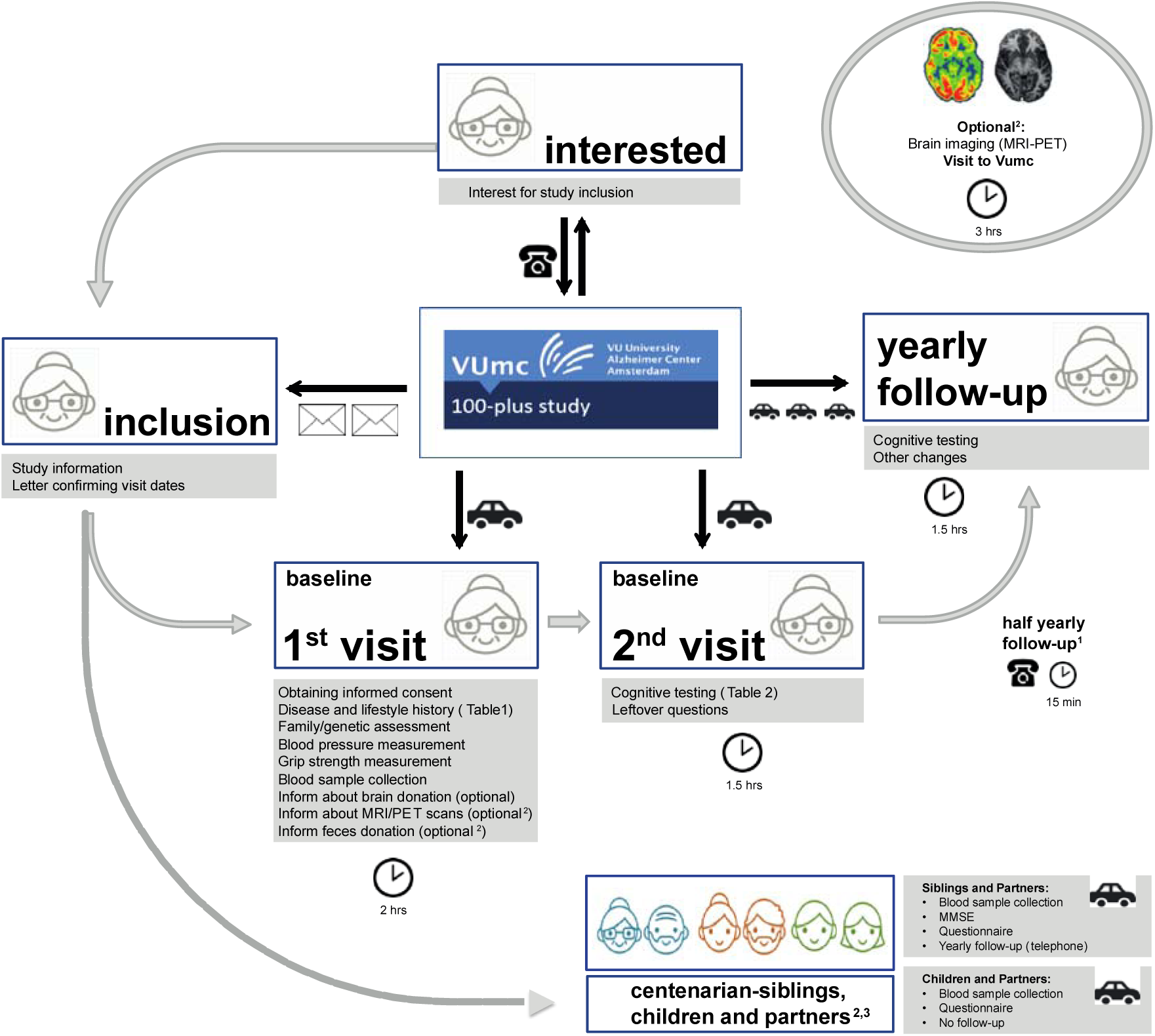
Diagram of visit procedures of 100-plus Study: ^1^Half yearly follow-up by telephone is performed for centenarians who agreed to brain donation. ^2^Collected in phase-2 of the 100-plus Study, started in September 2017. ^3^Data from centenarian-children and children in-laws will be obtained during the visit with the centenarian.

**Fig. 3:**
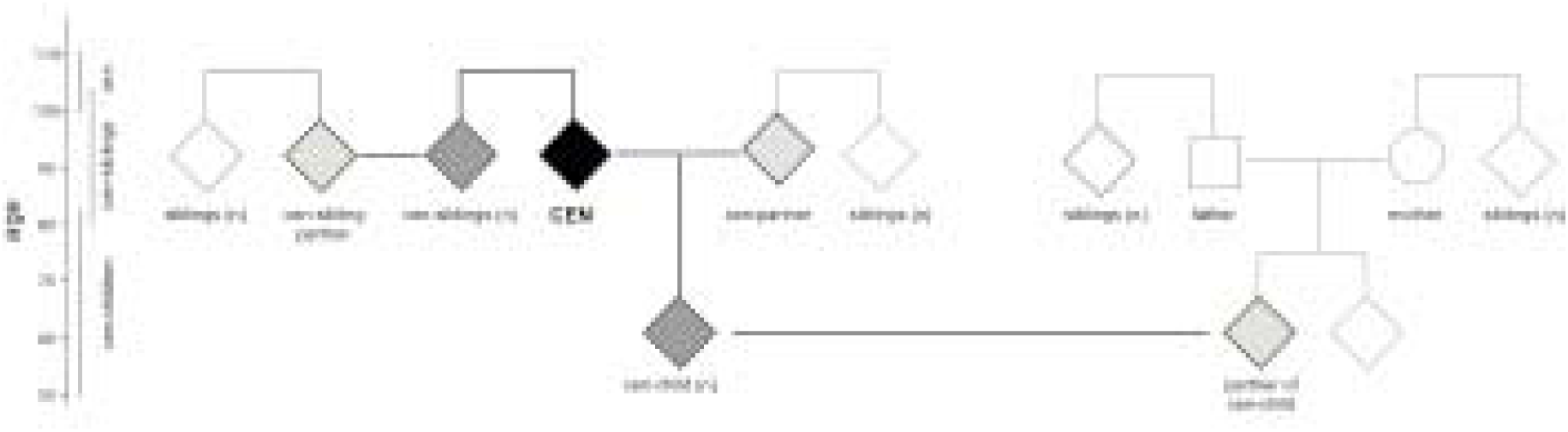
Data collection from centenarians and their family-members: In Phase-2 of the 100-plus Study (September 2017), we obtain blood-samples from centenarians (black), and when willing, their siblings, their children (dark grey) and their respective partners (light grey). We will inquire about longevity and incidence of dementia in relatives from the same generation as the centenarian (white). Square: male, circle: female, diamond: both genders are possible.

### Data collection

Centenarian presentation: During each visit, the researcher subjectively estimates the visual, hearing and mobility function as “good”, “moderate”, “poor” or “very poor”, according to the determinants listed in **Table 3**. We collect the following variables regarding centenarian presentation: the level of independence during activities of daily living <u>(ADL)</u> using the Barthel Index [27], an estimation of the total hours of care/assistance needed per week; category housing situation, (independent-dependent); grip strength, systolic and diastolic blood pressure; heartbeat; and napping habits and sleep quality (Pittsburg Sleep Quality Index questionnaire [28]). We assess whether the centenarian suffers from symptoms of depression [29, 30] by administering the 15-item geriatric depression scale (GDS-15). We ask about recent weight loss, current weight and length and active infections.

**Table 3:** Categorization of vision, hearing and mobility ability. Vision and hearing abilities were estimated while participants used all available devices to support their vision and/or hearing.

|  | Vision | Hearing | Mobility |
| --- | --- | --- | --- |
| **Good** | Able to read newspapers and watch television | Able to have and follow a conversation in a group of people | Able to walk independently (with or without help of a walking stick or walker) |
| **Moderate** | Able to read large texts with large letters and watch television | Able to have a conversation with one person/questions do not have to be repeated | Able to walk with help of another person |
| **Poor** | Not able to watch television/vision problems cause some difficulties in ADL | Limited ability to have a conversation with one person/questions need to be repeated multiple times | Able to move independently in a wheelchair |
| **Very poor** | Limited or complete loss of vision which causes severe difficulties in ADL | Not able to have a conversation with one person; this does not improve when speaking loud and clearly | Not able to move independently in a wheelchair |

#### Medical History

From each centenarian we request a General Practitioner (GP) summary report, which lists diagnosed conditions and prescribed medications. These conditions are categorized by a dedicated GP **(Table 4)**. After the centenarian died, we request a second synopsis from the GP, describing the medical proceedings until death. In a self-report medical history questionnaire, we inquire about blood pressure, heart disease; stroke (CVA) or TIA; tumors, head injuries, incontinence, dental condition, mental health problems, hospital visits, surgeries/anesthesia. We estimate BMI at midlife by recording self-reported weight and length at middle-age (∼50 years). For centenarian-females we inquire about age at menarche, onset of menopause, number of pregnancies and/or miscarriages.

**Table 4:** Categories of conditions analyzed in the GP medical files of 209 centenarians. Left column: when multiple conditions that belong to one condition-category are mentioned more than once in the GP report of a centenarian, they are counted as one. Right column: all conditions are counted separately, even though they belong to one condition-category. In aggregate, the percentages in the right column will exceed the percentage in the left column.

| Condition-category | Conditions |
| --- | --- |
| Fraction of centenarians with at least one mention of this condition in their GP report (%) | Fraction of centenarians with a at least one mention of these conditions in their GP report (%) |
| Cardiovascular disease (83.7%)<br>cardiovascular disease without hypertension (66.5%) | hypertension (48.8%); congestive heart failure (29.7%); cardiac dysrhythmia (23%); CVA/TIA (18.7%); angina pectoris (15.3%); myocardial infarction (8.1%); valvular heart disease (8.1%); thrombosis (6.2%); pacemaker (5.7%); aortic stenosis (2.9%); amputation leg (1.4%); coronary bypass (1%); hypercholesterolemia (1%); arterial disease (0.5%); arteritis temporalis (0.5%); atherosclerosis (0.5%); cerebrovascular insufficiency (0.5%); coronary sclerosis (0.5%); intermittent claudication (0.5%); orthostatic hypotension (0.5%); pericarditis (0.5%); |
| musculoskeletal (63.2%) | arthrosis (35.4%); fractures (34.4%); osteoporosis (14.8%); joint(s) replacement (11.5%); osteoarthritis (3.3%); hernia (1%); |
| vision (41.6%) | cataract (30.1%); macular (7.7%); glaucoma (3.8%); vision impairment (2.4%); |
| hearing (30.6%) | hearing impairment (30.6%); cholesteatoma (0.5%); sudden deafness (0.5%); |
| cancer (27.8%) | skin cancer (17.2%); breast cancer (4.3%); colon cancer (4.3%); prostate cancer (1.9%); uterus cancer (1.4%); bladder cancer (0.5%); cholesteatome (0.5%); palate cancer (0.5%); stomach cancer (0.5%); thyroid cancer (0.5%); vocal chord cancer (0.5%); |
| autoimmunology (22%) | diabetes (7.7%); rheumatoid arthritis (4.8%); hyperthyroidism (3.8%); hypothyroidism (3.3%); skin cancer (1.4%); asthma (1%); hypopituitarism (0.5%); thyroid enlargement (0.5%); thyroid removal (0.5%); |
| urology (21.5%) | UTI (7.2%); incontinence (5.7%); prostate hypertrophy (4.8%); hysterectomy (1.9%); uterine prolapse (1.9%); catheter (1%); prostate resection hypertrophy (1%); ovarian cysts (0.5%); |
| neurology/psychiatry (15.8%) | balance (3.3%); cognitive decline (2.9%); depression (2.4%); psychiatry (2.4%); epilepsy (1.9%); delirium (1.4%); insomnia (1%); Parkinson's (1%); dizziness (0.5%); migraine (0.5%); tremor (0.5%); WM atrophy (0.5%); |
| gastrointestinal (15.3%) | kidney failure (6.7%); gastric ulcer (1.9%); cholecystectomy (1.4%); diverticulosis (1.4%); gall stones (1.4%); kidney stones (1%); reflux esophagitis (1%); appendectomy (0.5%); intestinal polyps (0.5%); pancreatitis (0.5%); rectal prolapse (0.5%); sigmoid resection (0.5%); |
| lung disease (10.5%) | pneumonia (6.2%); COPD (2.4%); TBC (1.9%); Emphysema (0.5%); ulcer (0.5%); |
| other | erysipelas (1.4%); anemia (1%); herpes zoster (1%); other (1%); restless legs (1%); eye infection (0.5%); itching (0.5%); pes equinus (0.5%); vitamin B deficiency (0.5%); vitiligo (0.5%); |

#### Cognitive profiling

We objectively evaluate cognitive functioning using a comprehensive neuropsychological test battery that addresses memory, attention and/or concentration, pre-morbid intelligence, language, executive and visuo-spatial functions **(Table 2)**. To assess overall cognitive functioning we administer the Mini-Mental State Examination (MMSE) [26]. Geriatric sensory impairments such as bad eyesight or bad hearing complicated performance, which led to missing items. MMSE scores with different missing items cannot be directly compared, because the total obtainable score is different per centenarian. Therefore, we adjust scores using multiple imputation (**see ‘MMSE imputation’ in ESM.pdf**). In addition, at every visit the researcher subjectively estimated cognitive functioning of the centenarian (for procedures see ESM.pdf). During each research visit we ask an informant to fill in the Dutch version of the abbreviated form of the Informant Questionnaire on Cognitive

Decline (IQ-CODE) to indicate whether the centenarians experienced cognitive decline in the past ten years (or, in case of follow-up visits, during the past year) [31, 32].

#### Lifetime/demographic characteristics

To investigate the family genealogy and disease occurrence, we draw a pedigree including children, siblings, parents and grandparents, their (maiden) names, gender, birth years, age at death and cause of death, occurrence of dementia/cognitive decline. To determine socio-economic background (SEB) and socioeconomic status (SES) we inquire about the main occupation of the father and mother of the centenarian, the main occupation of the centenarian him/herself at adulthood and the main occupation of their partner(s). We inquire about the education level and the number of years education was followed. These were classified according to (I) ISCED 1997 [33] and according to Dutch 1971 census [34].

#### Lifetime habits

We address smoking habits and alcohol consumption **(see additional data)**. We administer the Cognitive Activity Questionnaire (CAQ) [35, 36] to investigate 1) cognitive stimulating experience during adult life (from childhood to 50 years) and 2) current cognitively stimulating experience.

#### Data-collection of first degree living centenarian-relatives and partners

For centenarian siblings and their partners, we administer the MMSE at the study inclusion visit and we record the genealogy at the level of the centenarian-generation. We will yearly monitor changes in physical well-being and in cognitive health using TICS-M and IQ-Code-N. We ask centenarian-children and partners to fill in an abbreviated version of the centenarian questionnaire; we record the genealogy of the centenarian-generation, no cognitive testing will be administered; we will not follow-up centenarian-children and their partners **(Table 1)**.

### Biomaterials

#### Biomaterial collection

During baseline visits, we collect a blood sample for DNA isolation, peripheral blood mononuclear cells (PBMCs), plasma, serum, and when consent is given for generation of induced pluripotent stem cells (iPSCs). DNA samples are currently used for APOE genotyping, GWAS, whole exome sequencing (WES) and Sanger sequencing. Furthermore, all centenarians are informed about the option for feces donation for gut microbiome analysis, PET-MRI or PET-CT brain scans for in vivo detection of amyloid beta presence and structural brain imaging. We also inform about the option of post-mortem brain donation. Brain autopsies are performed in collaboration with the Netherlands Brain Bank [37, 38]. For numbers of collected biomaterials thus far, please **see additional data** (ESM.pdf).

#### Data storage

OpenClinica open source software (version 3.1 onwards) is used for data management [39]. Biomaterials are stored in the biobank of the VUmc.

## COHORT DESCRIPTION

### Included centenarians

Between January 1^st^ 2013 and September 1^st^ 2018, 332 centenarians were included in the study of whom almost 30% (n=92) agreed to post mortem brain donation. Thus far, 58 centenarians have come to autopsy. Here we analyzed a first dataset, which was generated when the data collection from the first 300 centenarians was complete: on June 21^st^ 2017. At this time, 764 centenarians were approached for study-participation of which 300 (40%) met study-inclusion criteria and were included in the study **(Fig 4)**. For all cohort descriptives see **Table 5**. Additional data can be found in online supplementary material (ESM.pdf).

**Table 5:**
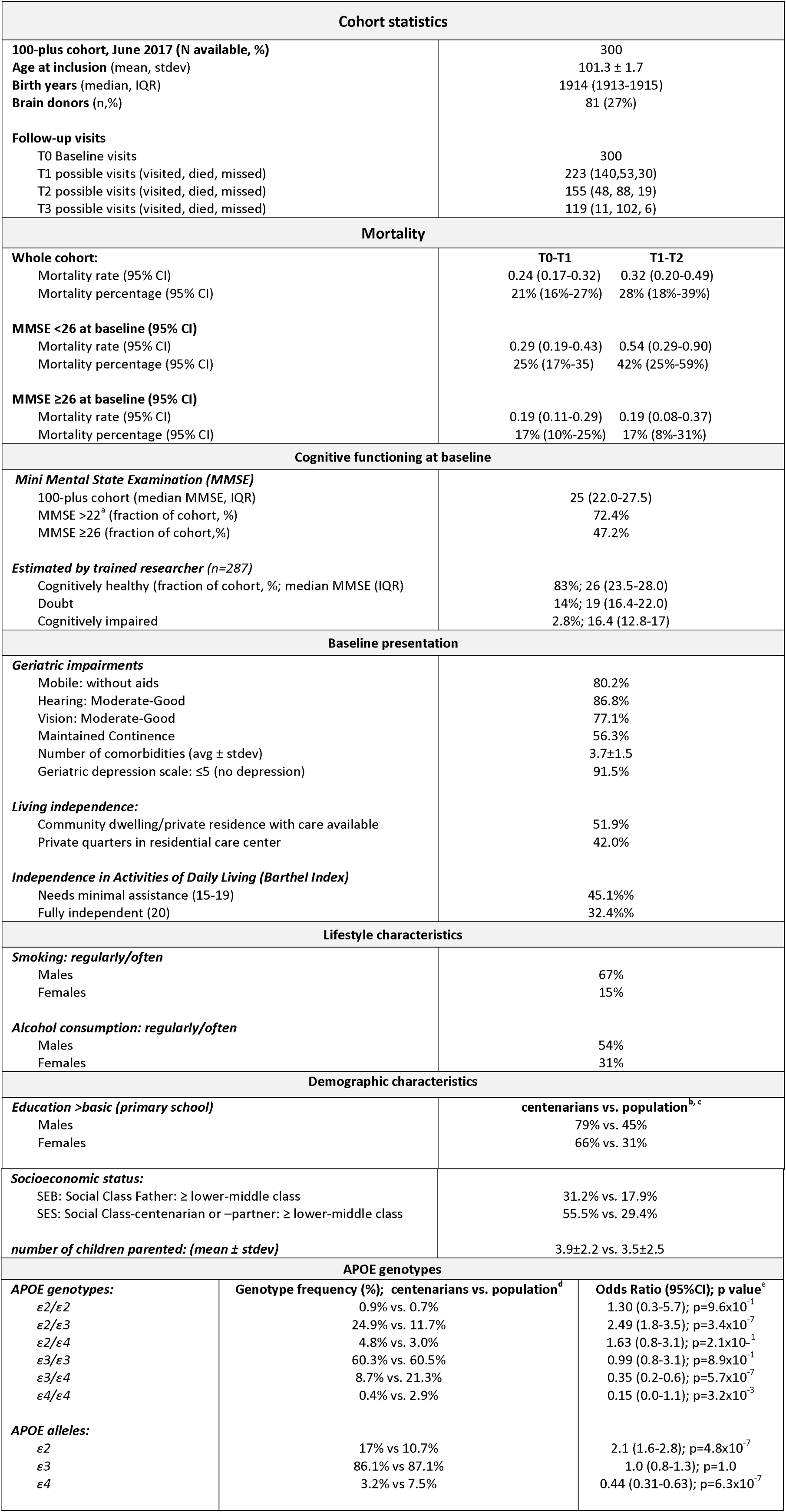
Descriptive statistics of 100-plus Study cohort. a: An MMSE >22 is the suggested cutoff score for cognitive health in elderly aged 97 years and above [118]. b: Centenarian education levels were compared with 54-61 years olds reported in the Dutch population in the 1971 census [40]; c: socio-economic background was compared with 2,815 individuals born between 1910-1915 from the Historical Sample of the Netherlands (HSN) [41]; d: APOE genotypes were compared with 2,233 ∼50-80-year olds from the Longitudinal Aging Study Amsterdam (LASA) [46]; e. P-values were calculated using a two-sided Fisher’s Exact test.

**Fig. 4:**
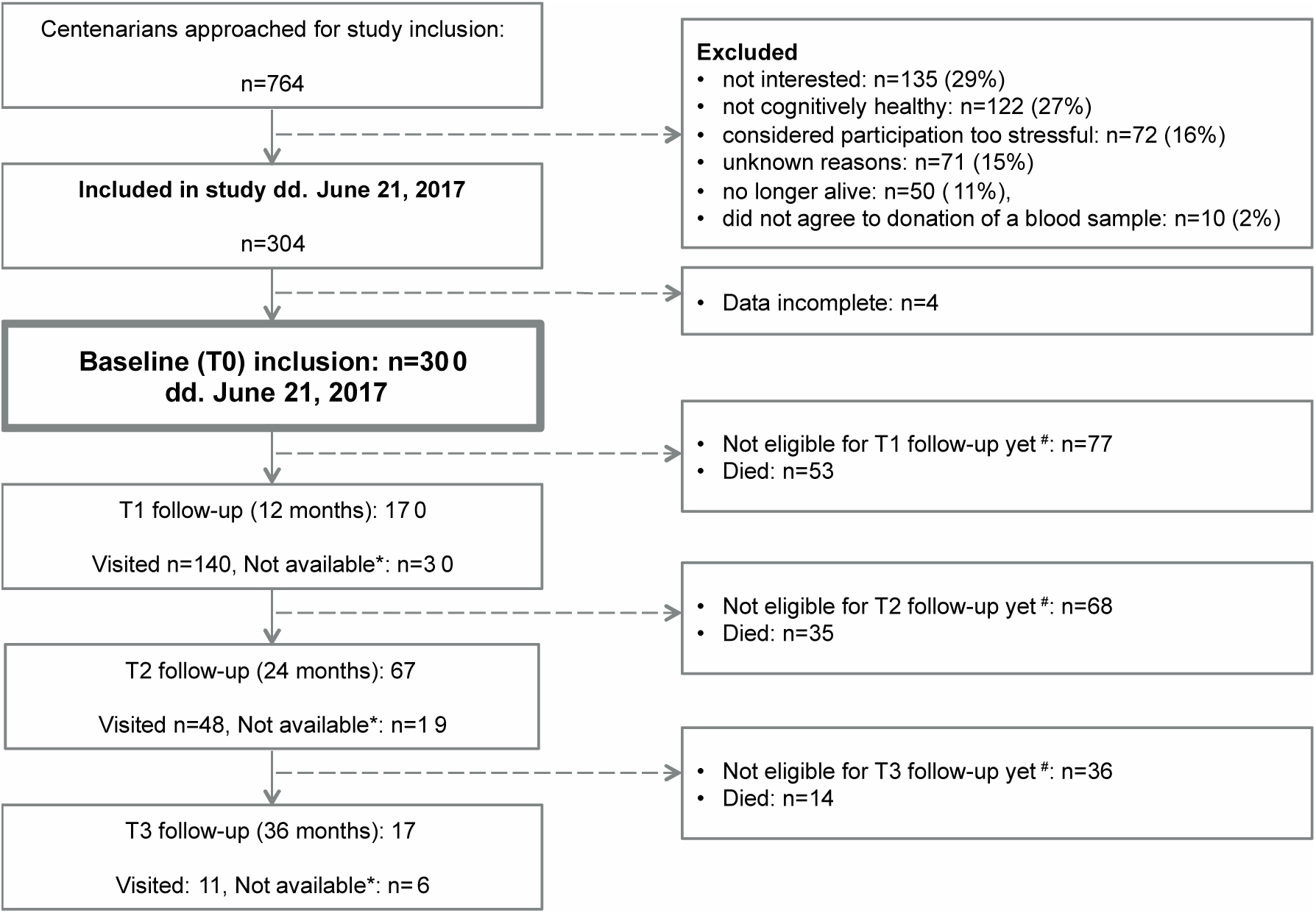
Flowchart of study inclusion: *Not available: centenarians were on vacation, not interested or too frail for a follow-up visit. When possible, follow-up was performed by telephone and/or informant questionnaires. In several cases, centenarians were available for follow-up one year later, such that this ‘unavailable’ group was formally kept in the study until death. ^#^Not eligible: centenarians were not yet included in the study long enough to be eligible for the next follow-up visit.

The mean age at inclusion of centenarians was 101.3±1.7 years (ESM.pdf **Fig S1A**). The majority of centenarians were born between 1910 and 1917 (ESM.pdf **Fig S1B**). Of the 300 centenarians in the cohort, 284 were born in a Dutch municipality, 6 were born in the Dutch East Indies, (a Dutch colony at the time), and 10 centenarians were born in other European countries. Centenarian birth-municipalities indicated that the catchment area is spread across the 11 Dutch provinces (ESM.pdf **Fig S2**).

### Presentation at baseline

Subjective researcher estimates of geriatric sensory impairments indicated that 87% of the centenarians had moderate-good hearing abilities (ESM.pdf **Fig S3A**), that 77% of the centenarians had moderate-good vision (ESM.pdf **Fig S3B**), and that 80% of the centenarians were independently mobile (ESM.pdf **Fig S3C**). The majority (52%) of the centenarians in the cohort lived independently (i.e. community dwelling without assistance, or independent in a residence with available services), 42% lived in private quarters in a residential care center, while only 1.7% of the centenarians lived in a nursing home (ESM.pdf **Fig S3D**). Centenarians scored a median of 15 points (IQR: 12-18), on the Barthel index: 45% of the centenarians scored between 15-19, which indicates a need for minimum help with activities of daily living (ADL), while 32% scored 20 points which indicates they are fully independent in ADL (ESM.pdf **Fig S3E**). The centenarians in the cohort have no or very few symptoms of depression: they scored a median of 2 points on the 15-items version of the Geriatric Depression Scale (IQR: 1-3), and scores <5 indicate no evidence for depression [29] (ESM.pdf **Fig S3F**).

### Disease prevalence and multi-morbidities

At the time of the data freeze we received and analyzed GP reports from 209 centenarians. At baseline, centenarians were diagnosed with or had symptoms of on average 3.7±1.5 morbidities (ESM.pdf **Fig S3G**). Cardiovascular problems are the most common condition in centenarians (83.7% has at least one mention of a cardiovascular condition in their GP report). And hypertension is mentioned in the GP reports of almost half of all centenarians. Removing hypertension from the list of cardiovascular conditions still leaves 66.5% of the centenarians with at least one mention of a cardiovascular condition (**Table 4**). Musculoskeletal disease and hypertension were more prevalent in females (72% vs. 39% and 54% vs. 34%), while cardiovascular conditions were more prevalent in males (77% vs. 63%). Most aging-associated diseases were first mentioned in the GP report when the centenarian was >90 years, suggesting a seemingly high age at onset. As we cannot correct for methodological differences in data collection by GPs, we were not able to perform a systematic comparison with disease incidence statistics from prospective cohort studies (for further explanation see ‘age at disease onset’ analysis in ESM.pdf).

### Cognitive function (Mini Mental State Examination, MMSE)

At cohort inclusion, the average raw MMSE score was 23.9±4.4 points. We adjusted for missing items due to hearing or vision impairments, which allowed us to directly compare MMSE scores between centenarians (see Methods). At study inclusion the average adjusted MMSE score of the 100-plus Study cohort was 24.3 ±4.23 points (median score: 25, IQR: 22.0-27.5) (**Fig 5A**). For 287 centenarians, a trained researcher estimated cognitive health. The large majority (83%) of the centenarians was subjectively estimated to be cognitively healthy, and this group scored a median of 26 points on the MMSE (IQR: 23.5-28). This was significantly higher than the median MMSE score of 19 (IQR: 16.4-22) by the 41 centenarians for whom cognitive health was “doubted” (p=4×10^-3^, two-tailed t-test with unequal variance), and the median MMSE score of 8 centenarians who were estimated to have “probable cognitive impairment” was 16.4, (IQR: 12.8-17) (**Fig 5B**).

**Fig. 5:**
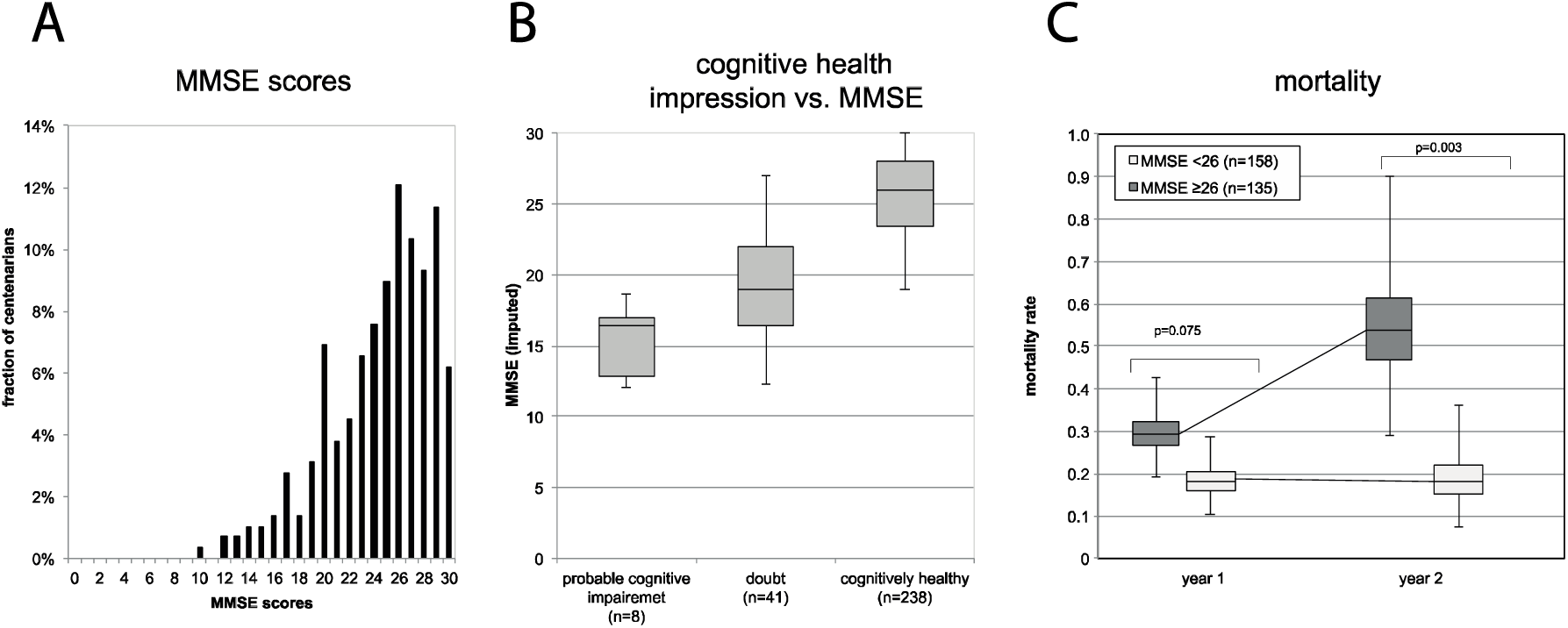
Overall cognitive functioning (Mini-Mental State Examination): **A.** Mini-Mental State Examination (MMSE) scores. **B.** Researcher impression of cognitive health at first visit, compared to MMSE score. **C.** Mortality rate of centenarians with high and low performance on the MMSE.

### MMSE and mortality rates

The mortality percentage (presented per annual-year) underestimates the mortality at extreme ages, such that we prefer presenting the instant mortality rates (presented per life-year); for rationale and calculation procedures see **ESM.pdf**). Within the group of 293 participants for which a baseline MMSE was available, there were 67 deaths that occurred before a next planned visit: the planning of a next visit was used to confirm which centenarians were still alive and who had died. There were 41 confirmed deaths that occurred before a planned first-year follow-up visit, and 174 centenarians were confirmed alive at the time of their first-year follow-up visit. The overall mortality rate in the first year after inclusion was 0.24 deaths per life-year (95%CI: 0.17-0.32); which relates to a mortality percentage of 21% per annual year (95%CI: 16%-27%). Specifically, the 106 centenarians who scored ≥26 on the MMSE at baseline had a mortality rate of 0.19 deaths per life-year (95%CI: 0.11-0.29), while the 109 centenarians with baseline MMSE scores <26 had a mortality rate of 0.29 deaths per annual-year (95%CI: 0.19-0.43) (p=0.075). Of the 91 centenarians who were eligible for a second follow-up visit, there were 20 confirmed deaths before this visit, and 71 were confirmed alive at the time of this visit. Therefore, in the second year after baseline, the mortality rate increased to 0.32 deaths per life-year (95%CI: 0.20-0.49); which relates to a mortality percentage of 28% per annual-year (95%CI: 18%-39%). Specifically, the mortality rate of the centenarians who scored ≥26 points at baseline remained at a low 0.19 deaths per life-year (95%CI: 0.08-0.37), while the mortality rate of centenarians who scored <26 points increased to 0.54 deaths per life-year (95%CI: 0.29-0.90) (p=3.0.×10^-3^) (**Fig 5C**). Mortality rates and related mortality percentages are presented in **Table 4.**

### Education

We retrospectively compared centenarian-education levels with 55-59 year-olds as reported in the Dutch population in the 1971 census [40], these individuals were from the same birth cohort as the centenarians (1912-1916). Both centenarian-males and females attained significantly higher levels of education compared to their birth cohort in the 1971 census [34] (p<1×10^-5^, Mann-Whitney U test) (ESM.pdf **Fig S4A**). Specifically, 79% of the centenarian males and 66% of the centenarian females attained more than basic education (primary school or less), compared to respectively 45% and 31% of the males and females in their birth cohort (**Table 5**). Workers and self-employed persons with little education were overrepresented in the ∼20% non-responders in the 1971 census, suggesting that this is a conservative estimate of the differences [34].

### Socio-economic background and status

Based on paternal professions, centenarian-socio-economic background (SEB) was compared to 2,815 individuals born between 1910-1915 from the Historical Sample of the Netherlands (HSN) [41] (p<1×10^-5^, Mann-Whitney U test) (ESM.pdf **Fig S4B-left**). Centenarian-fathers were 3-fold more likely to have an elite-upper middle class occupation and >3-fold less likely to be an unskilled worker compared to their birth cohort. Based on the professions of the 219 centenarian-males and centenarian-female-partners, centenarians themselves attained a significantly higher SES than the 408 males from the HSN sample born between 1910-1919 (p<1×10^-5^, Mann-Whitney U test). Centenarians were >4-fold more likely to be elite-upper middle class, >2-fold more likely to be farmers, and >3-fold less likely to be unskilled or farm workers (ESM.pdf **Fig S4B-right**). There was no difference between the socio-economic status (SES) attained as adults of 81 male centenarians and the male-partners of 138 female-centenarians (p=0.22, Mann-Whitney U test).

### Smoking behavior and alcohol consumption

Retrospective comparison of smoking behavior suggests that centenarians smoked less than a representative sample of Dutch individuals born between 1909-1923, as indicated in a 1958 survey [42, 43]. Of the centenarian-males, 67% indicated to have smoked regularly or often during an extended period in their lives, while 91% of the birth cohort males reported to smoke in 1958. Of the centenarian females, 15% indicated to have smoked regularly or often while 32% of the birth cohort females smoked. Alcohol consumption was common: only 11% of the centenarian-males and 22% of the centenarian-females indicated to never consume alcohol, similar to 14% of male-abstainers and 21.8% female-abstainers among the birth cohorts in the 1958 survey [42], whereas 54% of the centenarian-males and 31% of the centenarian-females indicated to consume alcohol regularly or often.

### Marriage and children

Centenarians had on average 3.9±2.2 children, which was more than the average 3.5±2.5 children from 860 Dutch parents born between 1910-1915 [44] (p=0.03, Mann-Whitney U) (ESM.pdf **Fig S4C**). However, we cannot exclude that lifestyle differences (i.e. religious or regional customs) might confound this increased fertility. Overall, 91% of the centenarians was ever-married, and 86% had one or more children. Of the centenarian females, 16.5% remained childless (36/219), similar to the 16% childless females born between 1915-1919 [45]. Five males in the cohort (6%) remained childless (birth cohort data not available [45]).

### APOE allele frequency

APOE was genotyped for 266 centenarians (ESM.pdf Fig S5). We observed that the centenarians were >2-fold more likely to carry an *APOE*-ε2 allele than 2,233 Dutch population controls aged 60-80 years [46]. Specifically, centenarians are 2.5-fold more likely to be genotyped APOE-ε2/ε3 (Table 5). In contrast, centenarians are >2-fold *less* likely to carry an *APOE*-ε4 allele compared to the Dutch population; specifically, centenarians are 2.8-fold less likely to be genotyped *APOE*-ε3/ε4 and 6.7-fold less likely to be genotyped *APOE*-ε4/ε4. The allele frequency of the *APOE* ε3 allele was identical for both cohorts.

## DISCUSSION

Here, we present the 100-plus Study cohort of cognitively healthy centenarians based on the first 300 centenarians included in the 100-plus Study.

### On average, the centenarians in the 100-plus Study cohort have a high performance on the MMSE; the large majority is independent and retained hearing and vision

Our inclusion criteria of *“self-reported cognitive health, which is confirmed by an informant”* sorted centenarians with a relatively high level of overall cognitive functioning. The cohort scored an average raw (unimputed) MMSE score of 23.9±4.4 points, which is considerably higher than average MMSE scores of representative centenarian populations (16.2 ± 8.8 points, Georgia Centenarian Study [47]; 18.7 ± 7.4, centenarians from central Italy (Rome and surroundings) [48]; 17.7 ± 8.3 centenarians from Northern Italy [49]). The overall cognitive performance of the 100-plus cohort participants is similar to *“community-dwelling cognitively healthy centenarians”* from the Georgia centenarian Study [50], and *“cognitively healthy”* Japanese centenarians, who respectively scored a mean of 24.8 points and 22.3±3.32 points on the MMSE [51].

Next to their retained cognitive functioning, the large majority of the centenarians had moderate-good hearing and vision abilities, they were independently mobile, they enjoyed a relatively high level of independence in activities of daily living (ADL), and had no or few symptoms of depression. Centenarians were either community dwelling or lived independently in a residence or in a care center with available services. Together, these findings echo that the 100-plus Study cohort is not a representative population of Dutch centenarians, rather, it represents a high-performing, independent sub-selection of Dutch centenarians.

### Cognitive performance is associated with mortality

The high cognitive and overall performance of our cohort was related with a two-fold reduced mortality relative to the centenarians in the general population. The first year after baseline, the mortality rate of the centenarians in our cohort was 0.24 deaths per life-year (translating to a mortality percentage of 21% per annual-year), which is ∼2-fold lower than the overall Dutch centenarian population with a mortality rate of almost 0.5 deaths per life-year (mortality percentage: 40% per annual-year) [52]. In the second year after baseline the overall mortality rate increased to 0.32 deaths per life-year (mortality percentage: 28% per annual-year); specifically, centenarians with high cognitive functioning retained a low mortality rate of 0.19 deaths per life-year (17% per annual-year), while centenarians with cognitive decline had a mortality rate of 0.54 deaths per life-year (42% per annual-year). These findings confirm that we have succeeded in selecting the healthiest of the centenarians, as we assume that these will be maximally enriched with protective (genetic) factors. Our results further confirm that there is an overlapping etiology of maintained cognitive and overall health, and cognitive functioning, which might be employed to predict overall decline and mortality [53, 54]. The longitudinal set-up of our study will allow us to monitor changes in cognition in combination with other factors of overall health that occur between baseline and death to identify to which extent centenarians escaped or delayed cognitive impairment.

### The 100-plus cohort is 2-fold enriched with males

The fraction of centenarian-males is 27%, twice the fraction of males (14.4%) in the total Dutch centenarian population on January 1^st^ 2017 [55]. Indeed, since dementia-prevalence in centenarian populations is consistently lower in males (∼40%) than in females (∼60%) [56, 57], we had on forehand expected that our inclusion criteria of *“self-reported cognitive health, which is confirmed by an informant”* might sort relatively more centenarian males than females. Based on the fraction of centenarian males in the population (14.3% in 2017) and lower dementia prevalence in males (40% vs. 60%), we estimated that the fraction of males in the 100-plus Study cohort should be approximately 20%. This suggests that dementia incidence does not fully explain the excess of males in our cohort, which leaves room for, for example, the influence of a participation bias and a better general well-being of centenarian males [58].

### Disease in male and female centenarians

Despite the cognitive health of the centenarians in the 100-plus Study cohort, they were diagnosed with on average four morbidities at baseline. Previous studies have shown that females are more prone to develop chronic nonfatal conditions such as dementia, arthritis and osteoporosis [59], while males are more likely to develop fatal conditions, such as cardiovascular disease and cancer [60, 61]. In agreement with these studies, we found that the females in the 100-plus Study cohort had a higher prevalence of musculoskeletal diseases and hypertension while males had a higher prevalence of heart disease and CVA/TIAs.

### The 100-plus Study cohort is equally depleted with the APOE-ε4 allele compared to other centenarian cohorts, but it is strongly enriched with the neuroprotective APOE-ε2 allele

The 100-plus Study cohort was 2.3-fold less likely to carry the APOE-ε4 AD-risk allele compared to their birth cohort at 60-80-years (OR=0.44, p=6.3×10^-7^) [46]. This is in complete concordance with the depletion observed in a meta-analysis of 2,776 (mostly Caucasian) centenarians and 12,000 controls (OR=0.43, p<1×10^-3^) [62]. It is well established that carrying one or two *APOE-ε*4 alleles is associated with respectively a 3-5 and 10-30-fold *increased* risk of developing AD. Depletion of the *APOE-ε*4 allele in centenarians supports an association of the ε4 allele with mortality at younger ages: carriers dropped out of the population during aging [63], leading to a depletion of the *APOE-ε*4 allele in centenarians that is consistent across studies.

On the other hand, carrying one protective *APOE-ε2* allele is associated with a 2-fold decreased lifetime risk of developing AD [64, 65]. But the large centenarian meta-analysis indicated that the protective aspect of the *APOE-ε2* allele does not extend to an enrichment in centenarians (OR=1.08, p=0.66), although a weak enrichment of the *APOE-ε2/ε3* genotype was observed (OR=1.4 p=1.7×10^-2^) [62]. In contrast, we observed that the APOE-ε2 allele was strongly enriched in cognitively healthy centenarians from the 100-plus Study cohort compared to their birth cohort at 60-80-years: centenarians were 2.1-fold more likely to carry the *APOE-ε2* allele (p=4.8×10^-7^), and 2.5-fold more likely to have an *APOE-ε2/ε3* genotype (p=3.4×10^-7^). This confirms previous suggestive findings in a cohort of Italian centenarians who were free of dementia or any other major age-related conditions, which had a similar enrichment of the *APOE-ε2* allele [66].

We speculate that this enrichment of the *APOE-ε2* allele is not a consequence of our selection of extreme ages, but that it reflects our selection of individuals with retained (cognitive) health until extreme ages.

The specific enrichment of the *APOE-ε2* in the centenarians with high cognitive performance suggests that the etiology for reaching 100 years with maintained (cognitive) health may be distinct from the etiology of reaching 100 years in general. Our results indicate that while searching for (genetic) factors that maintain cognitive health, the *APOE* genotype should be taken into account.

### Centenarians came, on average, from higher socio-economic classes and had higher levels of education

On average, centenarians came from a higher socio-economic background than their birth cohort. A high fraction of centenarian-fathers were farmers, mostly on their own farm, a common occupation in the Netherlands during the early twentieth century. As adults, centenarians attained a higher socio-economic status and they had more children compared to their birth cohort. Both male- and female-centenarians attained higher levels of education than the males and females from their birth cohorts. These findings reflect the selective survival advantage of individuals from the higher/middle socioeconomic classes and farmers, during the majority of the twentieth century in the Netherlands [67]. Together, this is in agreement with results from several centenarian studies, which showed that socioeconomic background, educational attainment, and adult socioeconomic status influenced the chance to become a centenarian [68]. Likewise, having children associates with an increased chance of reaching extreme ages, likely due to the involvement of children in the care for their aged parent [69].

### Alcohol consumption of centenarians was similar to birth cohort peers, and they smoked -but less

Two-thirds of the centenarian males and 15% of the centenarian females indicated to have smoked regularly or often during an extended period in their life. This was less than their birth cohort peers, of whom almost all males and a third of the females smoked [42]. Alcohol consumption of the centenarians was similar to their birth cohorts. These results are partly in agreement with lifestyle behaviours from the American Ashkenazi Jewish centenarians, whose alcohol consumption and smoking behaviour was not different from the general population [70].

We note that comparisons of lifestyle habits such as alcohol consumption and smoking rely on recall of habits they had several decades ago, which may introduce recall bias. For this reason, we focused on investigating lifestyle factors that are manifest for a longer period during a lifetime. Habits that may be more variable throughout life, such as dietary or exercise habits, might be more difficult to recall and we chose to refrain from investigating these. Despite limitations, the collected statistics can be applied in within-cohort analyses, such that they add to the rich phenotypic data available for this cohort.

## Conclusions

The 100-plus Study cohort represents cognitively healthy Dutch centenarians. Compared to their birth cohort peers, centenarians from this cohort attained significantly higher levels of education, were from a higher socioeconomic background, attained higher socioeconomic status, and they had more children, all of which confirms previous findings that these factors are associated with the chance of reaching 100 years in cognitive health. The combined contributions of these features, which are often concentrated within families, and the enrichment with the genetically heritable APOE-ε2 allele, will most likely explain a considerable proportion of the high heritability of reaching 100 years in maintained cognitive health. However, these features do not apply to all centenarians, and only a third of the cohort carries the APOE-ε2 allele. This suggests that additional protective factors may account for the cohort phenotype.

With the recent developments in biotechnology novel findings regarding the physiology of exceptional longevity and cognitive function are emerging [71, 72]. To advance such findings, the availability of blood and brain tissues of extensively phenotyped centenarians with the best possible aging-outcome and their family members, provides the opportunity to acquire insights in the molecular constellations associated with the long-term maintenance of cognitive health. Ultimately, with this cohort we aim to contribute to the generation of novel hypotheses regarding the generation of novel therapeutic targets that offer resilience to cognitive decline.

## CONFLICT OF INTEREST

The authors declare that they have no conflict of interest.

## ETHICS APPROVAL AND CONSENT TO PARTICIPATE

The Medical Ethical Committee of the VU University Medical Center has approved the 100-plus Study (METc-VUmc registration number: 2016.440). All procedures performed in studies involving human participants were in accordance with the ethical standards of the institutional and/or national research committee and with the 1964 Helsinki declaration and its later amendments or comparable ethical standards.

## INFORMED CONSENT

Informed consent was obtained from all individual participants included in the study.

## BOX: Study Rationale

The design of an intervention for neurodegenerative diseases requires not only the understanding of the neurodegenerative processes involved, but also a deep comprehension of the processes that *maintain* cognitive health during ageing. Although increasing age is the strongest predictive factor for cognitive decline and dementia, some people live to be over 110 years in great mental health. A Dutch woman, Hendrikje van Andel-Schipper (1890-2005), reached the age of 115 with full cognitive abilities [3] and showed that *it is possible to reach extreme ages without any symptoms of cognitive decline.* Her remarkable case became the source of inspiration for the initiation of the 100-plus Study at the VUmc Alzheimer Center in 2013. To investigate the physiology of her extended cognitive health, it is necessary to compare her clinical characteristics with those from other individuals with the same extraordinary combination of phenotypes: extremely old and cognitively healthy. Below, we provide a rationale for researching protective factors against cognitive decline in cognitively healthy centenarians, based on the mortality rate and dementia incidence in their birth cohort during its process of aging.

The number of centenarians in the Netherlands is growing quickly: on January 1^st^ 2013 there were 1940 centenarians in the Netherlands, which grew to 2,225 centenarians on January 1^st^ 2017, and this number is expected to rise to 5,000 by 2035 [55]. Of the individuals born between 1910-1915, approximately 1:160 (0.6%) have reached ages ≥100 years [73]. Since 25-30% of all centenarians are estimated to be free of symptoms of cognitive decline [74-77], becoming a centenarian with retained cognitive health was reserved for only 0.2-0.3% of the 1910-1915 birth generation.

Almost all participants of the 100-plus Study cohort were born in the Netherlands just before or during WWI (1914-1917), in which the Netherlands was neutral. The 20^th^ century in the Netherlands was further characterized by a depression in the thirties, WWII between 1940-1945, a post-war period typified by a rebuilding phase in the 1950s and a continuous increase in prosperity, health care improvements and technological developments. According to the Human Mortality Database [73] males born in the Netherlands between 1910-1915 had a mean lifespan of 58.7 years and females had a mean lifespan of 66.1 years.

Here we describe the 1910-1915 birth cohort by their mortality rates and dementia incidence from birth to >100 years (**Box-Fig**). For this, we prefer presenting the instant mortality rate over the mortality percentage, because, while estimates are similar at younger ages, the mortality percentage underestimates the mortality at extreme ages (for further explanation **see Mortality estimations in ESM.pdf**). For ages 0-60 years, we represent mortality rate by age from individuals born in 1912, and for ages >60 years we represent mortality rates using combined statistics from the 1910-1915 birth cohorts.

(**A**) In 1912, 170,000 babies were born, and they were exposed to a mortality rate of close to 0.10 deaths life-year (∼10% per annual-year) during their first year of life, reflecting prevalent childhood diseases. (**B**) In 1918, the Spanish Flu made >40,000 casualties in the Netherlands [78], which was especially lethal among 20-40 year olds [79]. The Spanish Flu increased mortality among the 1912-born six-year olds by 2-fold. (**C**) At age 10, most childhood diseases were overcome and the mortality rate reached a stable ∼0.001-0.002 deaths per life-year (∼0.10%-0.20% per annual-year), caused by incidental deaths (i.e. fatal accidents, drowning etc. (**D**) When the 1912 birth cohort was 28 years, the onset of WWII in 1940 led to a peak in male mortality. (**E**) The ensuing Hunger Winter in 1945 led to a second mortality peak when the 1912 birth cohort was 33 years old, more so in males than in females. (**F**) When the cohort was 40 years old, the mortality resulting from natural decline rose above the rate of incidental deaths, and increased in the log scale according to Gompertz Law (1825) [80]. (**G**) During natural decline of the 1912 birth cohort, the males had a higher mortality rate than females, and this mortality gender gap ultimately resulted in a 1:7 male/female ratio at age 100 [55]. Approximately 70% of the mortality gender gap in these cohorts can be explained by the difference in smoking behavior between males and females [81]: an estimated 91% of all males born in 1912 smoked while only 30% of the females smoked, which ultimately led to a relative increased incidence of fatal smoking-related diseases in males. The remainder of the mortality gender gap may be explained by biological or environmental differences between males and females [82]. (**H**) At 100 years old, the 1912 cohort has reduced to ±1,000 persons and the instant mortality rate for both males and females is at 0.5 deaths per life-year (which translates to a mortality percentage of ∼40% per annual-year). (**I**) Individuals from the 1910-1915 birth cohorts were exposed to an increasingincidence of overall dementia from age 60 years onwards, of which the greatest proportion was (**J**) Alzheimer’s Dementia (AD) [1]. At approximately 100 years, the instant incidence of dementia reaches 0.5 cases per dementia-free year, which translates to a dementia proportion of ∼40% per annual-year. At this age, the dementia incidence surpasses the mortality rate per year, suggesting that after turning 100 years, a centenarian is exposed to greater odds of developing dementia than to die [6]. (**K**) If dementia incidence after 100 years continues to increase exponentially, following the Gompertz law of natural decline [80], then a conservative estimation of dementia incidence (by concentrating mortality on incident dementia cases) suggests that all individuals who reach 108-110 years would have to be demented. (**K_2_**) In contrast, reports of individuals who are older than 110 years indicate that the majority of such individuals has, in fact, retained their cognitive health [3-5]. Therefore, it is likely that the incidence of dementia decelerates or even declines at extreme ages [6]. Although the slope of the incidence rate suggested by Corrada et al. (red dots in Fig) is slightly smaller compared to the extrapolated incidence (dashed line), there currently is no clear evidence for this deceleration between 90 and 100 years [2, 83], it is most likely that this deceleration becomes evident somewhere after 100 years. This is consistent with findings in super-centenarians by Andersen et al. [5], who demonstrated the progressive compression of both disability and morbidity (in 6 diseases including dementia) with survival beyond 100 years. Furthermore, in a recent study based on data from 3,836 centenarians in Italy, Barbi et al found that mortality decelerates, and even plateaus, above age 105 [84]. Together, this suggests that factors that preserve (cognitive) health may be progressively enriched for during healthy aging [7], providing a window of opportunity to search for such protective factors in a population of healthy (super-) centenarians.

**Box-Figure.**
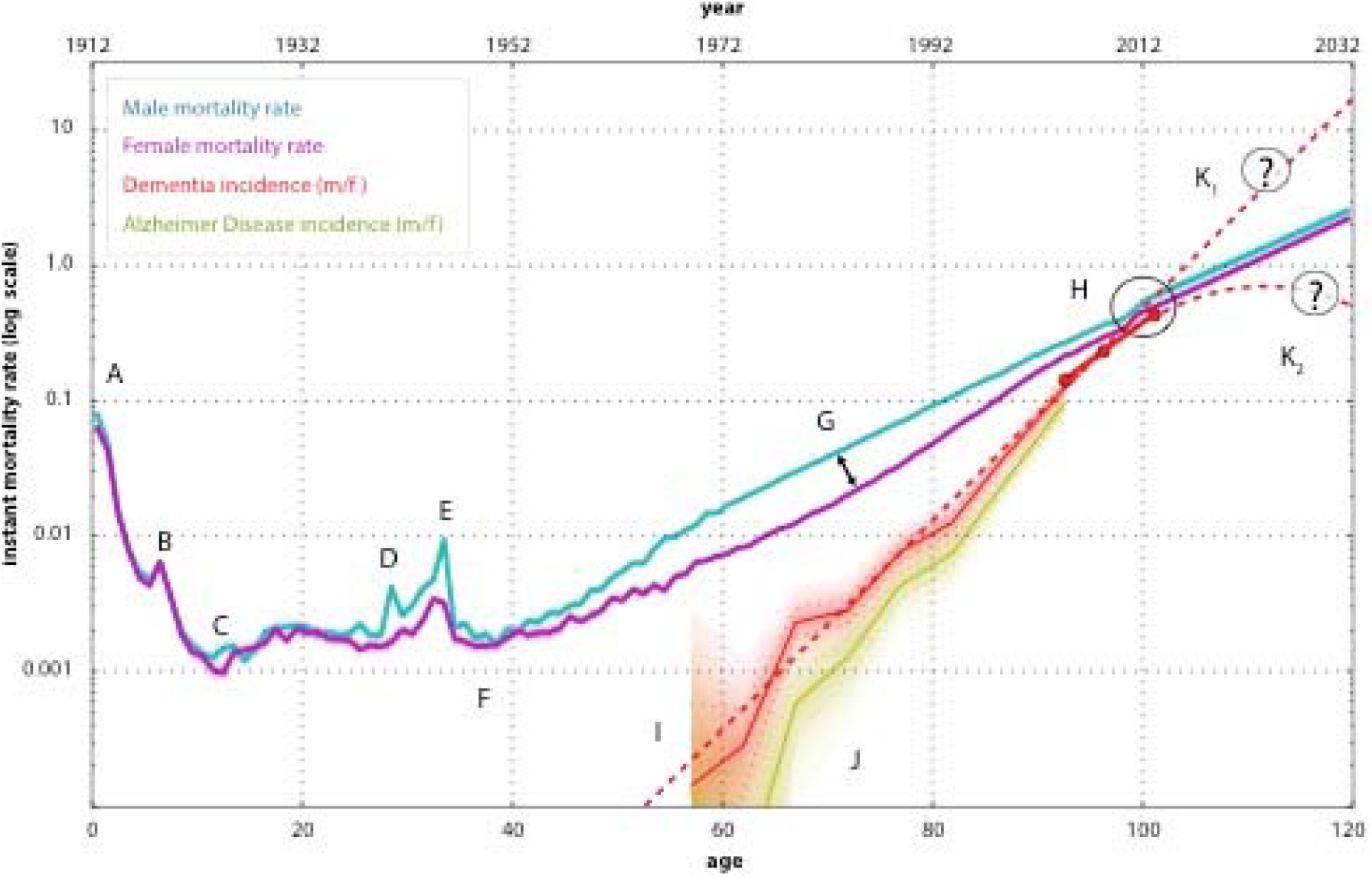
Instant mortality and dementia rates in centenarian birth cohort. Blue line: Male mortality. Shades of blue represent confidence intervals (CI) on the mortality rate by 10-percentile increments [85]; For ages 0-59 years we used only the mortality statistics of individuals born in 1912 (as to avoid blurring specific mortality peaks), and for ages 60-100 years we combined statistics of the 1910-1915 birth cohorts, which reduced CIs. Mortality after age 100 years was extrapolated in accordance with the Gompertz’ law of mortality [86]. **Purple line:** Female mortality with CIs [85]. **Red line:** Median incidence of overall dementia with CIs for age groups 55-59, 60-64, 65-65, 70-74, 75-79, 80-84, 85-89 years [1]. To define mean age per age-group, we assumed that the ages of the individuals that constituted each age-group were distributed according to associated mortality statistics. Red dots: Dementia incidence for age groups: 90-94 years (mean age 92.7), 95-99 years (mean age 96.4) and 100+ (mean age 101.3) [2]. Green line: Alzheimer’s Disease (AD) incidence with CIs [1]. **Dashed red line:** extrapolation of dementia incidence according to its exponential increase. To extrapolate dementia incidence, we fitted a Gompertz curve on available dementia incidence data [1, 2]. For the reported age ranges we compared the estimated dementia incidence with the reported incidence through a binomial distribution. This resulted in a log-likelihood, which was optimized (see ESM.pdf for mortality calculations).

AUTHORS’ CONTRIBUTIONS
HH conceived and designed the study and wrote the manuscript; NB helped obtain ethical approval for the study and with the management of communication with participants and their proxies; TD is dedicated GP and analyzed GP reports, KP, EW, KS, LT, DB and SR visited the centenarians and helped design/improve the study as it evolved; SS advised on the design of the neuropsychological testing battery; FvP provided expert knowledge on occupation classification and the demographics of the 1910-1920 birth cohorts, HMH contributed substantive support for study set-up; MH helped with the study design and manages the Open Clinica database for data storage; PS contributed substantive support for the study and aided with obtaining necessary funding. All authors read and approved the final manuscript.

## ACKNOWLEDGEMENTS

We are grateful for the collaborative efforts of all participating centenarians and their family members and/or relations. We also want to acknowledge the many people who contributed and continue to contribute to the study. The students involved in recruiting and visiting the centenarians: Lieke Jansma, Lieve Steins Bisschop, Sanne Koole, Sanne Hofman, Anna Kamsteeg, Saiedah Wekker, Matteo Neumann, and Marlous Jansen. The personnel who facilitated data collection and storage in available biobanks: Michiel Kooreman, Annemieke Rozemuller, Jeroen Hoozemans, Hans Gille, Charlotte Teunissen. The persons who advised on the collection procedures of specific data: Quinten Waisfisz, Eus van Someren, Ted Koene, Bart van Berckel, Wiesje van der Flier, Vivi Heine, Erik Sistermans, Cornelia van Duijn. We thank Marcel Reinders and Sven van der Lee for critically reviewing the manuscript before submission.

## Notes

**FUNDING:** This work was supported by Stichting Alzheimer Nederland (WE09.2014-03), Stichting Diorapthe (VSM 14 04 14 02) and Stichting VUmc Fonds.

## REFRENCES

1. Lobo A, Lopez-Anton R, Santabarbara J, de-la-Camara C, Ventura T, Quintanilla MA et al. Incidence and lifetime risk of dementia and Alzheimer’s disease in a Southern European population. Acta psychiatrica Scandinavica. 2011;124(5):372–83. doi:10.1111/j.1600-0447.2011.01754.x.

2. Corrada MM, Brookmeyer R, Paganini-Hill A, Berlau D, Kawas CH. Dementia incidence continues to increase with age in the oldest old: the 90+ study. Annals of neurology. 2010;67(1):114–21. doi:10.1002/ana.21915.

3. den Dunnen WF, Brouwer WH, Bijlard E, Kamphuis J, van Linschoten K, Eggens-Meijer E et al. No disease in the brain of a 115-year-old woman. Neurobiol Aging. 2008;29(8):1127–32. doi:10.1016/j.neurobiolaging.2008.04.010.

4. Jeune B, Robine J-M, Young R, Desjardins B, Skytthe A, Vaupel JW. Jeanne Calment and her successors. Biographical notes on the longest living humans. 2010:285–323. doi:10.1007/978-3-642-11520-2_16.

5. Andersen SL, Sebastiani P, Dworkis DA, Feldman L, Perls TT. Health Span Approximates Life Span Among Many Supercentenarians: Compression of Morbidity at the Approximate Limit of Life Span. The Journals of Gerontology Series A: Biological Sciences and Medical Sciences. 2012;67A(4):395–405. doi:10.1093/gerona/glr223.

6. Robine JM, Jagger C. What Do We Know About the Cognitive Status of Supercentenarians? In: Christen Y, editor. Research and Perspectives in Longevity; Longevity and Frailty. Springer; 2003. p. 145–52.

7. Barzilai N, Atzmon G, Derby CA, Bauman JM, Lipton RB. A genotype of exceptional longevity is associated with preservation of cognitive function. Neurology. 2006;67(12):2170–5. doi:10.1212/01.wnl.0000249116.50854.65.

8. Barral S, Cosentino S, Costa R, Andersen SL, Christensen K, Eckfeldt JH et al. Exceptional memory performance in the Long Life Family Study. Neurobiol Aging. 2013;34(11):2445–8. doi:10.1016/j.neurobiolaging.2013.05.002.

9. Cosentino S, Schupf N, Christensen K, Andersen SL, Newman A, Mayeux R. Reduced prevalence of cognitive impairment in families with exceptional longevity. JAMA neurology. 2013;70(7):867–74. doi:10.1001/jamaneurol.2013.1959.

10. Haworth CM, Wright MJ, Martin NW, Martin NG, Boomsma DI, Bartels M et al. A twin study of the genetics of high cognitive ability selected from 11,000 twin pairs in six studies from four countries. Behavior genetics. 2009;39(4):359–70. doi:10.1007/s10519-009-9262-3.

11. Petrill SA, Saudino K, Cherny SS, Emde RN, Fulker DW, Hewitt JK et al. Exploring the genetic and environmental etiology of high general cognitive ability in fourteen-to thirty-six-month-old twins. Child development. 1998;69(1):68–74.

12. Petrill SA, Kovas Y, Hart SA, Thompson LA, Plomin R. The genetic and environmental etiology of high math performance in 10-year-old twins. Behavior genetics. 2009;39(4):371–9. doi:10.1007/s10519-009-9258-z.

13. Plomin R, Haworth CM. Genetics of high cognitive abilities. Behavior genetics. 2009;39(4):347–9. doi:10.1007/s10519-009-9277-9.

14. Perls TT, Wilmoth J, Levenson R, Drinkwater M, Cohen M, Bogan H et al. Life-long sustained mortality advantage of siblings of centenarians. Proceedings of the National Academy of Sciences of the United States of America. 2002;99(12):8442–7. doi:10.1073/pnas.122587599.

15. Adams ER, Nolan VG, Andersen SL, Perls TT, Terry DF. Centenarian offspring: start healthier and stay healthier. Journal of the American Geriatrics Society. 2008;56(11):2089–92. doi:10.1111/j.1532-5415.2008.01949.x.

16. Mostafavi H, Berisa T, Day FR, Perry JRB, Przeworski M, Pickrell JK. Identifying genetic variants that affect viability in large cohorts. Plos Biol. 2017;15(9):e2002458. doi:10.1371/journal.pbio.2002458.

17. Sebastiani P, Andersen SL, McIntosh AI, Nussbaum L, Stevenson MD, Pierce L et al. Familial Risk for Exceptional Longevity. North American actuarial journal: NAAJ. 2016;20(1):57–64. doi:10.1080/10920277.2015.1061946.

18. Harper AR, Nayee S, Topol EJ. Protective alleles and modifier variants in human health and disease. Nature reviews Genetics. 2015;16(12):689–701. doi:10.1038/nrg4017.

19. Bucci L, Ostan R, Cevenini E, Pini E, Scurti M, Vitale G et al. Centenarians’ offspring as a model of healthy aging: a reappraisal of the data on Italian subjects and a comprehensive overview. Aging (Albany NY). 2016;8(3):510–9. doi:10.18632/aging.100912.

20. Atzmon G, Schechter C, Greiner W, Davidson D, Rennert G, Barzilai N. Clinical phenotype of families with longevity. Journal of the American Geriatrics Society. 2004;52(2):274–7.

21. Silverman JM, Smith CJ, Marin DB, Birstein S, Mare M, Mohs RC et al. Identifying families with likely genetic protective factors against Alzheimer disease. American journal of human genetics. 1999;64(3):832–8. doi:10.1086/302280.

22. Liu JZ, Erlich Y, Pickrell JK. Case–control association mapping by proxy using family history of disease. Nature genetics. 2017;49(3):325–31. doi:10.1038/ng.3766.

23. Jansen I, Savage J, Watanabe K, Bryois J, Williams D, Steinberg S et al. Genetic meta-analysis identifies 10 novel loci and functional pathways for Alzheimer’s disease risk. bioRxiv. 2018;prepublication. doi:10.1101/258533.

24. Marioni RE, Harris SE, Zhang Q, McRae AF, Hagenaars SP, Hill WD et al. GWAS on family history of Alzheimer’s disease. Translational psychiatry. 2018;8(1). doi:10.1038/s41398-018-0150-6.

25. Bucci L, Ostan R, Cevenini E, Pini E, Scurti M, Vitale G et al. Centenarians’ offspring as a model of healthy aging: a reappraisal of the data on Italian subjects and a comprehensive overview. Aging. 2016;8(3):510–9. doi:10.18632/aging.100912.

26. Folstein MF, Folstein SE, McHugh PR. “Mini-mental state”. A practical method for grading the cognitive state of patients for the clinician. Journal of psychiatric research. 1975;12(3):189–98.

27. Mahoney FI, Barthel DW. Functional Evaluation: The Barthel Index. Maryland state medical journal. 1965;14:61–5.

28. Buysse DJ, Reynolds CF, 3rd, Monk TH, Berman SR, Kupfer DJ. The Pittsburgh Sleep Quality Index: a new instrument for psychiatric practice and research. Psychiatry research. 1989;28(2):193–213.

29. Yesavage JA, Sheikh JI. 9/Geriatric Depression Scale (GDS). Clinical Gerontologist. 1986;5(1-2):165–73. doi:10.1300/J018v05n01_09.

30. Bleeker JAC, de Winter FML, Cornelissen E. Geriatric Depression Scale. 1985.

31. Jorm AF, Jacomb PA. The Informant Questionnaire on Cognitive Decline in the Elderly (IQCODE): socio-demographic correlates, reliability, validity and some norms. Psychological medicine. 1989;19(4):1015–22.

32. de Jonghe JF, Schmand B, Ooms ME, Ribbe MW. [Abbreviated form of the Informant Questionnaire on cognitive decline in the elderly]. Tijdschrift voor gerontologie en geriatrie. 1997;28(5):224–9.

33. UNESCO. International Standard Classification of Education 1997. http://www.unesco.org/education/information/nfsunesco/doc/isced_1997.htm. 1997.

34. Vliegen JM, de Jong U, Wesselingh AA, van der Kley P, CBS, SISWO. Education in the Netherlands. Monografiëën volkstelling 1971. The Hague s-Gravenhage: Centraal Bureau voor de Statistiek: Staatsuitgeverij; 1981.

35. Wilson R, Barnes L, Bennett D. Assessment of lifetime participation in cognitively stimulating activities. Journal of clinical and experimental neuropsychology. 2003;25(5):634–42. doi:10.1076/jcen.25.5.634.14572.

36. Ossenkoppele R. Cognitieve Activities Questionnaire. 2013.

37. Netherlands Brain Bank. http://www.brainbank.nl/.

38. Ravid R, Swaab DF. The Netherlands brain bank--a clinico-pathological link in aging and dementia research. Journal of neural transmission Supplementum. 1993;39:143–53.

39. Open Clinica. Copyright © OpenClinica LLC and collaborators, Waltham, MA, USA http://www.OpenClinica.com.

40. Statistics Netherlands. 14e Algemene Volkstelling annex woningtelling 28 februari 1971. The Hague: Centraal Bureau voor de Statistiek; 1980. p. 40.

41. Mandemakers K. Historical sample of the Netherlands HSN. Historical Social Research. 2001;26(4):179–90.

42. Gadourek I. Riskante gewoonten en zorg voor eigen welzijn (Hazardous habits and human well-being). Groningen: J.B. Wolters; 1963.

43. Nagelhout G. Trendpublicatie percentage rokers. Percentage rokers in de Nederlandse bevolking 1958 – 2012 2013.

44. Van Poppel F, Reher D, Sanz-Gimeno A, Sanchez-Dominguez M, Beekink E. Mortality decline and reproductive change during the Dutch demographic transition. Demographic Research. 2012;27:299–338. doi:10.4054/DemRes.2012.27.11.

45. Dykstra PA. Childless Old Age. In: Uhlenberg P, editor. International Handbook of Population Aging. Springer; 2009. p. 671–90.

46. Huisman M, Poppelaars J, van der Horst M, Beekman AT, Brug J, van Tilburg TG et al. Cohort profile: the Longitudinal Aging Study Amsterdam. International journal of epidemiology. 2011;40(4):868–76. doi:10.1093/ije/dyq219.

47. Miller LS, Mitchell MB, Woodard JL, Davey A, Martin P, Poon LW et al. Cognitive Performance in Centenarians and the Oldest Old: Norms from the Georgia Centenarian Study. Aging, Neuropsychology, and Cognition. 2010;17(5):575–90. doi:10.1080/13825585.2010.481355.

48. Bauco C, Borriello C, Cinti AM, Martella S, Zannino G, Rossetti C et al. Correlation between MMSE performance, age and education in centenarians. Archives of gerontology and geriatrics. 1998;26:23–6. doi:10.1016/s0167-4943(98)80004-6.

49. Baldelli MV, Salvioli G, Neri M, Pradelli JM. A survey of a centenarian population in Italy, focusing on self-sufficiency and cognition. Archives of gerontology and geriatrics. 1996;22 Suppl 1:345–54. doi:10.1016/0167-4943(96)86960-3.

50. Holtsberg PA, Poon LW, Noble CA, Martin P. Mini-Mental State Exam status of community-dwelling cognitively intact centenarians. International psychogeriatrics. 1995;7(3):417–27.

51. Inagaki H, Gondo Y, Hirose N, Masui Y, Kitagawa K, Arai Y et al. Cognitive Function in Japanese Centenarians according to the Mini-Mental State Examination. Dementia and geriatric cognitive disorders. 2009;28(1):6–12. doi:10.1159/000228713.

52. Central Bureau of Statistics: Prognosis period-life expectancy; gender and age, 2014-2060 [database on the Internet]. Netherlands Statistics. 2017. Available from: http://statline.cbs.nl/Statweb/publication/?DM=SLNL&PA=82690ned&D1=0&D2=a&D3=a&D4=0,16,l&VW=T. Accessed: 29 Aug 2017

53. Skoog J, Backman K, Ribbe M, Falk H, Gudmundsson P, Thorvaldsson V et al. A Longitudinal Study of the Mini-Mental State Examination in Late Nonagenarians and Its Relationship with Dementia, Mortality, and Education. Journal of the American Geriatrics Society. 2017;65(6):1296–300. doi:10.1111/jgs.14871.

54. Mossakowska M, Broczek K, Wieczorowska-Tobis K, Klich-Raczka A, Jonas M, Pawlik-Pachucka E et al. Cognitive performance and functional status are the major factors predicting survival of centenarians in Poland. The Journals of gerontology Series A, Biological sciences and medical sciences. 2014;69(10):1269–75. doi:10.1093/gerona/glu003.

55. Centraal Bureau voor de Statistiek. Aantal 100-plussers verdubbeld in 20 jaar. Webmagazine CBS 2017.

56. Yang Z, Slavin MJ, Sachdev PS. Dementia in the oldest old. Nature reviews Neurology. 2013;9(7):382–93. doi:10.1038/nrneurol.2013.105.

57. Corrada MM, Brookmeyer R, Berlau D, Paganini-Hill A, Kawas CH. Prevalence of dementia after age 90: results from the 90+ study. Neurology. 2008;71(5):337–43. doi:10.1212/01.wnl.0000310773.65918.cd.

58. Hazra NC, Dregan A, Jackson S, Gulliford MC. Differences in Health at Age 100 According to Sex: Population-Based Cohort Study of Centenarians Using Electronic Health Records. Journal of the American Geriatrics Society. 2015;63(7):1331–7. doi:10.1111/jgs.13484.

59. Pinn VW. Past and future: sex and gender in health research, the aging experience, and implications for musculoskeletal health. The Orthopedic clinics of North America. 2006;37(4):513–21. doi:10.1016/j.ocl.2006.09.006.

60. Wingard DL, Cohn BA, Kaplan GA, Cirillo PM, Cohen RD. Sex differentials in morbidity and mortality risks examined by age and cause in the same cohort. American journal of epidemiology. 1989;130(3):601–10.

61. Roman-Blas JA, Castaneda S, Largo R, Herrero-Beaumont G. Osteoarthritis associated with estrogen deficiency. Arthritis research & therapy. 2009;11(5):241. doi:10.1186/ar2791.

62. Garatachea N, Marin PJ, Santos-Lozano A, Sanchis-Gomar F, Emanuele E, Lucia A. The ApoE gene is related with exceptional longevity: a systematic review and meta-analysis. Rejuvenation research. 2015;18(1):3–13. doi:10.1089/rej.2014.1605.

63. Gerdes LU, Jeune B, Ranberg KA, Nybo H, Vaupel JW. Estimation of apolipoprotein E genotype-specific relative mortality risks from the distribution of genotypes in centenarians and middle-aged men: apolipoprotein E gene is a “frailty gene,” not a “longevity gene”. Genetic epidemiology. 2000;19(3):202–10. doi:10.1002/1098-2272(200010)19:3<202::AID-GEPI2>3.0.CO;2-Q.

64. Genin E, Hannequin D, Wallon D, Sleegers K, Hiltunen M, Combarros O et al. APOE and Alzheimer disease: a major gene with semi-dominant inheritance. Molecular psychiatry. 2011;16(9):903–7. doi:10.1038/mp.2011.52.

65. Farrer LA, Cupples LA, Haines JL, Hyman B, Kukull WA, Mayeux R et al. Effects of age, sex, and ethnicity on the association between apolipoprotein E genotype and Alzheimer disease. A meta-analysis. APOE and Alzheimer Disease Meta Analysis Consortium. JAMA: the journal of the American Medical Association. 1997;278(16):1349–56.

66. Garatachea N, Emanuele E, Calero M, Fuku N, Arai Y, Abe Y et al. ApoE gene and exceptional longevity: Insights from three independent cohorts. Experimental gerontology. 2014;53:16–23. doi:10.1016/j.exger.2014.02.004.

67. Kunst AE, Looman CW, Mackenbach JP. Socio-economic mortality differences in The Netherlands in 1950-1984: a regional study of cause-specific mortality. Social science & medicine. 1990;31(2):141–52.

68. Fors S, Lennartsson C, Lundberg O. Childhood living conditions, socioeconomic position in adulthood, and cognition in later life: exploring the associations. The Journals of gerontology Series B, Psychological sciences and social sciences. 2009;64(6):750–7. doi:10.1093/geronb/gbp029.

69. Modig K, Talback M, Torssander J, Ahlbom A. Payback time? Influence of having children on mortality in old age. Journal of epidemiology and community health. 2017;71(5):424–30. doi:10.1136/jech-2016-207857.

70. Rajpathak SN, Liu Y, Ben-David O, Reddy S, Atzmon G, Crandall J et al. Lifestyle Factors of People with Exceptional Longevity. Journal of the American Geriatrics Society. 2011;59(8):1509–12. doi:10.1111/j.1532-5415.2011.03498.x.

71. Zainabadi K. A brief history of modern aging research. Experimental gerontology. 2018;104:35–42. doi:10.1016/j.exger.2018.01.018.

72. Scheltens P, Blennow K, Breteler MM, de Strooper B, Frisoni GB, Salloway S et al. Alzheimer’s disease. Lancet. 2016;388(10043):505–17. doi:10.1016/S0140-6736(15)01124-1.

73. CBS/HMD. Unpublished data from the NIDI mortality database provided by Ewa Tabeau (NIDI) (1850-1949); Central Bureau of Statistics 1949-onwards (Data obtained through the Human Mortality Database, http://www.mortality.org, on 29-5-2017).

74. Poon LW, Woodard JL, Stephen Miller L, Green R, Gearing M, Davey A et al. Understanding dementia prevalence among centenarians. The Journals of gerontology Series A, Biological sciences and medical sciences. 2012;67(4):358–65. doi:10.1093/gerona/glr250.

75. Kliegel M, Moor C, Rott C. Cognitive status and development in the oldest old: a longitudinal analysis from the Heidelberg Centenarian Study. Archives of gerontology and geriatrics. 2004;39(2):143–56. doi:10.1016/j.archger.2004.02.004.

76. Andersen-Ranberg K, Vasegaard L, Jeune B. Dementia is not inevitable: a population-based study of Danish centenarians. The Journals of gerontology Series B, Psychological sciences and social sciences. 2001;56(3):P152–9.

77. Calvert JF, Jr., Hollander-Rodriguez J, Kaye J, Leahy M. Dementia-free survival among centenarians: an evidence-based review. The Journals of gerontology Series A, Biological sciences and medical sciences. 2006;61(9):951–6.

78. Ansart S, Pelat C, Boelle PY, Carrat F, Flahault A, Valleron AJ. Mortality burden of the 1918-1919 influenza pandemic in Europe. Influenza and other respiratory viruses. 2009;3(3):99–106. doi:10.1111/j.1750-2659.2009.00080.x.

79. Taubenberger JK, Morens DM. 1918 Influenza: the mother of all pandemics. Emerging infectious diseases. 2006;12(1):15–22. doi:10.3201/eid1201.050979.

80. Gompertz B. On the Nature of the Function Expressive of the Law of Human Mortality, and on a New Mode of Determining the Value of Life Contingencies. Philosophical Transactions of the Royal Society of London. 1825;115.:513–83.

81. Janssen F, van Poppel FWA. Roken veroorzaakte vroeger grote verschillen in levensverwachting. DEMOS: Bulletin over Bevolking en Samenleving. 2016:4–6.

82. Newman AB, Murabito JM. The epidemiology of longevity and exceptional survival. Epidemiologic reviews. 2013;35:181–97. doi:10.1093/epirev/mxs013.

83. James BD, Schneider JA. Increasing incidence of dementia in the oldest old: evidence and implications. Alzheimer’s research & therapy. 2010;2(3):9. doi:10.1186/alzrt32.

84. Barbi E, Lagona F, Marsili M, Vaupel JW, Wachter KW. The plateau of human mortality: Demography of longevity pioneers. Science. 2018;360(6396):1459–61. doi:10.1126/science.aat3119.

85. Central Bureau of Statistics: Life expectancy per birth cohort. [database on the Internet]. Netherlands Statistics. 2017. Available from: http://statline.cbs.nl/Statweb/dome/?TH=26190&PA=80333NED&LA=nl. Accessed: 11 Aug 2015

86. Gavrilov LA, Gavrilova NS. Mortality Measurement at Advanced Ages: A Study of the Social Security Administration Death Master File. North American actuarial journal: NAAJ. 2011;15(3):432–47.

87. Kok RM, Verhey FRJ. Gestandaardiseerde MMSE. Altrecht GGZ. 2002.

88. Nelson HE, O’Connell A. Dementia: the estimation of premorbid intelligence levels using the New Adult Reading Test. Cortex; a journal devoted to the study of the nervous system and behavior. 1978;14(2):234–44.

89. Schmand B, Lindeboom J, van Harskamp F. Nederlandse leestest voor volwassenen.. In: Bouma A, Mulder J, Lindeboom J, editors. Neuropsychologische Diagnostiek. Lisse: Swets & Zeitlinger; 1992. p. A45–A52.

90. Schmand B, Bakker D, Saan R, Louman J. De Nederlandse Leestest voor Volwassenen: een maat voor het premorbide intelligentieniveau. Gerontologie en Geriatrie. 1991;22:15–9.

91. de Jager CA, Budge MM, Clarke R. Utility of TICS-M for the assessment of cognitive function in older adults. International journal of geriatric psychiatry. 2003;18(4):318–24. doi:10.1002/gps.830.

92. Morris JC, Heyman A, Mohs RC, Hughes JP, van Belle G, Fillenbaum G et al. The Consortium to Establish a Registry for Alzheimer’s Disease (CERAD). Part I. Clinical and neuropsychological assessment of Alzheimer’s disease. Neurology. 1989;39(9):1159–65.

93. Lindeboom J, Schmand B, Tulner L, Walstra G, Jonker C. Visual association test to detect early dementia of the Alzheimer type. Journal of neurology, neurosurgery, and psychiatry. 2002;73(2):126–33.

94. Wilson BA, Cockburn J, Baddeley AD. The Rivermead behavioural memory test. 2nd ed., rev., enl. and redesigned. ed. Thames Valley Test Company; 1991.

95. Van Balen HGG, Groot Zwaaftink AJM. Rivermead Behavioural Memory Test. Nederlandse bewerking. In Thames Valley test Company, Reading. 1987.

96. Lindeboom J, Matto D. [Digit series and Knox cubes as concentration tests for elderly subjects]. Tijdschrift voor gerontologie en geriatrie. 1994;25(2):63–8.

97. Wechsler D. Wechsler Adult Intelligence Scale–Revised. San Antonio, TX: Psychological Corporation; 1981.

98. Wechsler D. WAIS-III administration and scoring manual. San Antonio, TX: Psychological Corporation; 1997.

99. Army Individual Test Battery. Manual of Directions and Scoring. 1944.

100. Reitan RM. Validity of the Trail Making Test as an indicator of organic brain damage. Perceptual and Motor Skills. 1958;8:271–6. doi:10.2466/PMS.8.7.271-276.

101. Van der Elst W, Van Boxtel MP, Van Breukelen GJ, Jolles J. Normative data for the Animal, Profession and Letter M Naming verbal fluency tests for Dutch speaking participants and the effects of age, education, and sex. Journal of the International Neuropsychological Society: JINS. 2006;12(1):80–9. doi:10.1017/S1355617706060115.

102. Lezak MD. Neuropsychological assessment. 2nd edition. ed.

103. Schmand B, Groenink SC, van den Dungen M. [Letter fluency: psychometric properties and Dutch normative data]. Tijdschrift voor gerontologie en geriatrie. 2008;39(2):64–76.

104. Snijders J, Luteijn F, van der Ploeg F, Verhage F. Handleiding Groninger intelligentie test. Lisse: Swets & Zeitlinger; 1983.

105. Benton AL, Hamsher KD. Multilingual Aphasia Examination. Iowa City, IA: AJA Associates; 1989.

106. Krabbendam L, Kalff AC. Handleiding Nederlandse vertaling BADS, (B.A. Wilson, N. Alderman, P.W. Burgess, H. Emslie en J.J. Evans). 1997.

107. Wilson BA, Evans JJ, Alderman N, Burgess PW, Emslie H. Behavioural assessment of the dysexecutive syndrome. Methodology of frontal and executive function. 1997:239–50.

108. Lindeboom J, Jonker C. Amsterdamse Dementie Screeningstest. Lisse, The Netherlands: Swets and Zeitlinger; 1988.

109. Luteijn F, van der Ploeg FAE. Groninger Intelligentie Test, Handleiding. 1982.

110. Roth M, Tym E, Mountjoy CQ, Huppert FA, Hendrie H, Verma S et al. CAMDEX. A standardised instrument for the diagnosis of mental disorder in the elderly with special reference to the early detection of dementia. The British journal of psychiatry: the journal of mental science. 1986;149:698–709.

111. Roth M, Huppert F, Tym E, Mountjoy CQ. CAMDEX-R boxed set: the revised cambridge examination for mental disorders of the elderly. 1998:81–8.

112. Shulman KI, Pushkar Gold D, Cohen CA, Zucchero CA. Clock-drawing and dementia in the community: A longitudinal study. International journal of geriatric psychiatry. 1993;8(6):487–96. doi:10.1002/gps.930080606.

113. Shulman KI. Clock-drawing: is it the ideal cognitive screening test? International journal of geriatric psychiatry. 2000;15(6):548–61.

114. Warrington EK, James M. The Visual Object and Space Battery Perception. 1991.

115. Collin C, Wade DT, Davies S, Horne V. The Barthel ADL Index: a reliability study. International disability studies. 1988;10(2):61–3.

116. de Haan R, Limburg M, Schuling J, Broeshart J, Jonkers L, van Zuylen P. [Clinimetric evaluation of the Barthel Index, a measure of limitations in dailly activities]. Nederlands tijdschrift voor geneeskunde. 1993;137(18):917–21.

117. Post MW, van Asbeck FW, van Dijk AJ, Schrijvers AJ. [Dutch interview version of the Barthel Index evaluated in patients with spinal cord injuries]. Nederlands tijdschrift voor geneeskunde. 1995;139(27):1376–80.

118. Kahle-Wrobleski K, Corrada MM, Li B, Kawas CH. Sensitivity and specificity of the minimental state examination for identifying dementia in the oldest-old: the 90+ study. Journal of the American Geriatrics Society. 2007;55(2):284–9. doi:10.1111/j.1532-5415.2007.01049.x.

